# Environment-dependent behavioral traits and experiential factors shape addiction vulnerability

**DOI:** 10.1101/2020.04.12.038141

**Authors:** Maxime Fouyssac, Mickaël Puaud, Eric Ducret, Lucia Marti-Prats, Nathalie Vanhille, Solène Ansquer, Xinxuan Zhang, Aude Belin-Rauscent, Chiara Giuliano, Jean-Luc Houeto, Barry. J Everitt, David Belin

**Affiliations:** Department of Psychology, University of Cambridge, Downing street, CB2 3EB, Cambridge, UK; Université de Poitiers, Faculté de Pharmacie, 6 Rue de la Milétrie, 86073 Poitiers, France; INSERM CIC-1402, CHU of Poitiers, 2 Rue de la Milétrie, 86021 Poitiers, France

**Keywords:** cocaine, alcohol, compulsivity, environmental enrichment, schedule-induced polydipsia

## Abstract

The transition from controlled drug use to drug addiction depends on an interaction between a vulnerable individual, their environment and a drug. Here we tested the hypothesis that conditions under which individuals live influence behavioral vulnerability traits and experiential factors operating in the drug taking environment to determine the vulnerability to addiction. The role of behavioural vulnerability traits in mediating the influence of housing conditions on the tendency to acquire cocaine self-administration was characterised in 48 rats housed in either an enriched (EE) or a standard (SE) environment. Then, the influence of these housing conditions on the individual vulnerability to develop addiction-like behaviour for cocaine or alcohol was measured in 72 EE or SE rats after several months of cocaine self-administration or intermittent alcohol drinking, respectively. The determining role of negative experiential factors in the drug taking context was further investigated in 48 SE rats that acquired alcohol drinking as a self-medication strategy. The environment influenced the acquisition of drug intake through its effect on behavioral markers of resilience to addiction. In contrast, the initiation of drug taking as a coping strategy or in a negative state occasioned by the contrast between enriched housing conditions and a relatively impoverished drug taking setting, facilitated the development of compulsive cocaine and alcohol intake. These data demonstrate that addiction vulnerability depends on environmentally determined experiential factors, suggesting that initiating drug use through negative reinforcement-based self-medication facilitates the development of addiction in vulnerable individuals.

**Significance Statement:** The factors that underlie an individual’s vulnerability to switch from controlled, recreational drug use to addiction are not well understood. We showed that in individuals housed in enriched conditions, the experience of drugs in the relative social and sensory impoverishment of the drug taking context, and the associated change in behavioural traits of resilience to addiction, exacerbate the vulnerability to develop compulsive drug intake. We further demonstrated that the acquisition of alcohol drinking as a mechanism to cope with distress increases the vulnerability to develop compulsive alcohol intake. Together these results demonstrate that experiential factors in the drug taking context shape the vulnerability to addiction.

## Introduction

Ten to forty percent of individuals recreationally using drugs are estimated eventually to develop the compulsive drug seeking and taking (1) that characterise drug addiction (2). It has long been considered that this individual vulnerability to transition from controlled to compulsive drug taking stems from the interaction between genetically determined behavioral traits, the nature of the neurodevelopmental environment and the context in which the drug is taken (3). However, the evident limitations of longitudinal studies in humans, which cannot readily control environmental and experiential factors (4, 5), and the inherent limitations of preclinical animal models (6) have made understanding the nature of these interactions and their underlying mechanisms extremely difficult.

The discovery of individual differences in the vulnerability to develop addiction-like behavior (7, 8), measured by operationalising behaviorally in rats three key features of the diagnosis of the disorder in the DSM (9), has helped to identify factors that mediate the effects of gene x drug interactions on the tendency to take drugs and the subsequent vulnerability or resilience to addiction (10). For example, high locomotor reactivity to novelty, suggested to operationalize the human trait of sensation seeking, has been shown to predict an increased tendency to self-administer stimulants (11), and also to be to be a marker of resilience to the transition to cocaine addiction (7, 12). This is consistent with evidence that sensation seeking in humans is related to recreational drug use, but not to stimulant addiction (13). Other behavioral traits (10), such as anxiety, impulsivity and novelty preference, have been shown specifically to predict loss of control over cocaine intake (14) or the transition to cocaine- or alcohol-addiction like behavior in rats, mice and in humans (7, 12, 13, 15).

How these behavioral factors of vulnerability are influenced by, or interact with, the living or drug use environment to shape the propensity to engage in drug taking is not yet well understood. Similarly, it has not been established whether experiential determinants of drug taking, such as seeking sensory or affective enhancement, or coping with distress (4, 16), independently mediate the vulnerability to transition from controlled to compulsive drug taking.

The impact of an individual’s environment on the vulnerability to addiction has to date been considered to be both unidimensional and unidirectional: impoverished environmental conditions such as those experienced by individuals in low socio-economic groups, or, inferentially, by rodents kept in impoverished housing conditions, or social isolation (SI), are considered to promote addiction (17). However, recent epidemiological data bring this into question (18). Experimentally, seemingly paradoxical observations include those showing that rats raised in an enriched environment (EE) are less likely to self-administer amphetamine (19), methylphenidate (20) or cocaine (21), yet are actually more sensitive to the reinforcing and motivational effects of cocaine than rats raised in social isolation (SI) (22). EE rats show lower self-administration titration rates than SI rats under fixed ratio schedules of reinforcement (17, 19), and also freely drink more alcohol under two-bottle choice conditions than SI rats (23).

These observations are consistent with the epidemiological evidence of a relative increase in premature drug-related deaths among individuals from high socio-economic populations (18), who also drink more often and consume higher quantities of alcohol than those from lower socio-economic backgrounds, even though the latter seem to experience more negative consequences of drug use (24). It may therefore be hypothesized that experiential factors related to drug use, rather than living conditions per se, are important determinants of the vulnerability to addiction across environmental conditions, as also indicated by the rise in drug-related deaths observed in very wealthy individuals (25, 26).

Here we tested the hypothesis that housing conditions and experiential factors operating in the drug taking environment interact with behavioral vulnerability traits to determine the likelihood of making the transition from controlled to compulsive drug use. The influence of differential housing conditions on the vulnerability to develop addiction-like behavior was investigated by comparing EE rats to rats raised in pairs in a social environment (SE) in standard cages, thereby controlling for any confounding effect that social isolation may independently exert on the psychological and behavioral mechanisms under investigation (27–29).

## Results

### Environmental enrichment decreases the propensity to acquire cocaine self-administration in addiction-resilient rats

We first causally tested the influence of different housing conditions, i.e. EE vs SE (n=24 each) on behavioral traits that are related to personality factors relevant to addiction in humans, namely anxiety (14), sensation seeking (7, 11, 30), boredom susceptibility (12), reward sensitivity (31) and sign-tracking (32) (**Fig. S1**), that we propose to reflect a multidimensional behavioural profile in the rat (**see SOM**). EE abolished the drug use proneness/addiction-resilience trait of high locomotor reactivity to novelty (HR), decreased the anxiety-associated behaviors of head dipping on the elevated plus-maze (EPM); EE also disrupted the asymmetric conditioned approach behavior usually displayed by sign-trackers towards a food CS, in line with previous evidence that EE impairs the attribution of incentive salience to food-paired cues (33) (**Fig. 1 and Fig. S2, S3**). EE did not influence each trait independently, but it shaped their relative multidimensional configuration in three distinct multidimensional behavioural profiles identified at the population level by cluster analysis (**Fig. 1B**). In particular EE increased the probability of displaying a multidimensional behavioural profile driven by sweetness and novelty preference (model 3), as opposed to that driven by stress reactivity (11, 34–36) (model 1) that is preferentially displayed by SE rats (**Fig. 1B**).

**Figure1.**
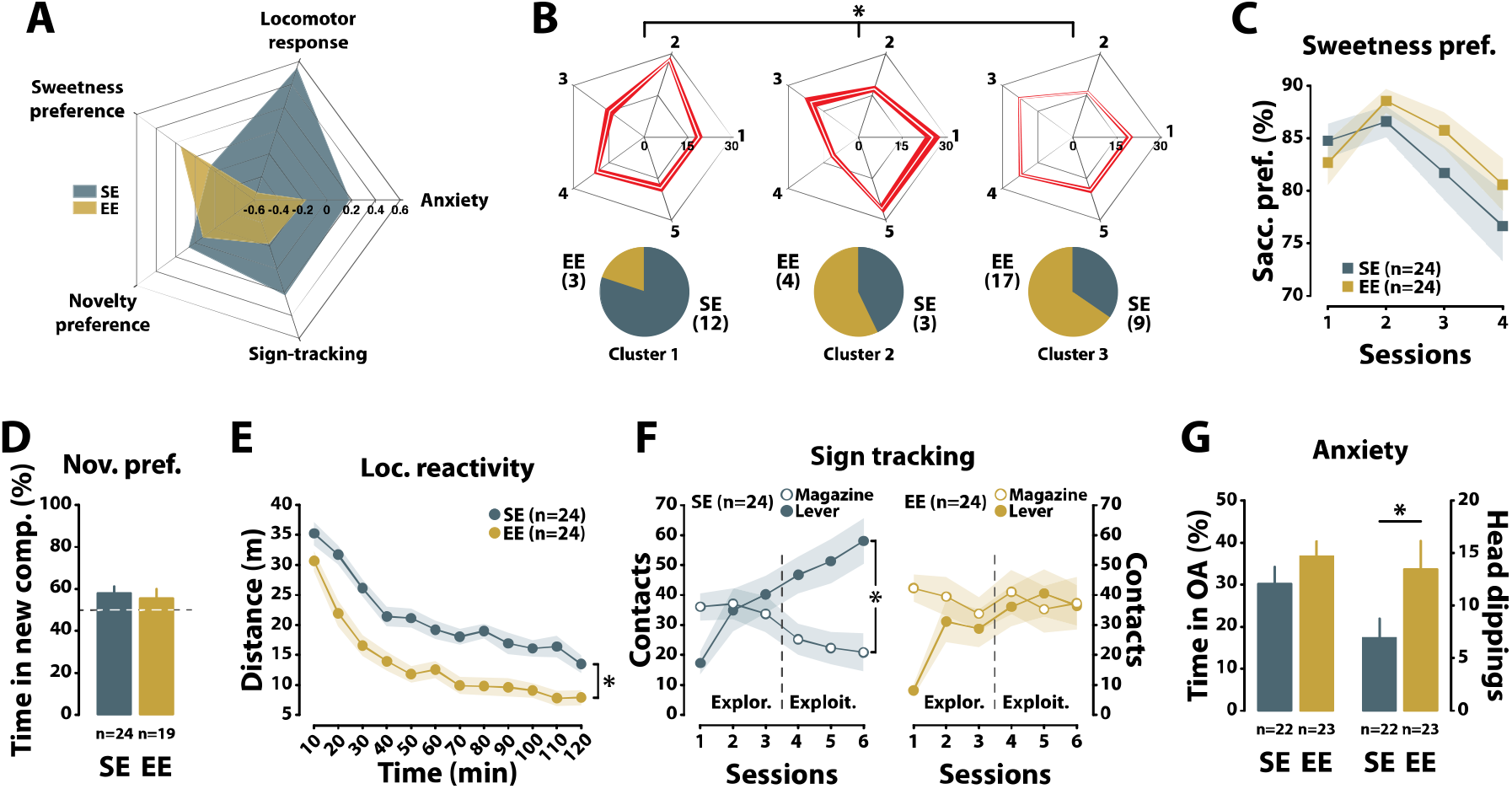
Environmental enrichment differentially shapes the behavioral traits of vulnerability and resilience to addiction in rats. Enrichment of the housing environment (EE) exacerbates the contribution of sweetness preference to a multidimensional behavioral model of personality diminishing the contribution of anxiety, sign-tracking and reactivity to novelty **(A and B)**. These behavioral traits differed in their contribution to the three distinct, environment-dependent [EE vs SE: Chi2=8.43, p<0.05], multidimensional behavioural profiles identified from a cluster analysis [cluster x trait interaction: F_8,180_=14.373, p<0.0001, ηp^2^<=0.39], thereby suggesting a complex multidimensional interaction between the environmental conditions and behavioral traits of vulnerability versus resilience to addiction. Quantitatively, EE did not influence the sweetness preference of rats [main effect of group: F_1,46_<1; session: F_3,138_=11.538, p<0.0001, ηp^2^<=0.20 and group x session interaction: F_3,138_=1.7583, p>0.05] **(C)** nor their preference for novelty [main effect of group: F_1,41_=2.3124, p>0.05] **(D)**. However, EE abolished the behavioral trait of resilience to addiction, namely high locomotor response to novelty (HR), in that EE rats showed a marked decrease in their locomotor reactivity to novelty as compared to SE rats [main effect of group: F_1,46_=19.274, p<0.0001, ηp^2^<=0.30; time: F_11,506_=102.16, p<0.0001, ηp^2^<=0.69; group × time interaction: F(11,506)=1.5757, p>0.05] **(E)**. Similarly, EE prevented the development of asymmetrical approach behavior, biased towards the CS, characteristically displayed by sign tracker rats raised in a SE during the exploitation phase of an AutoShaping task [SE, left panel: phase x approach response: F_1,23_=24.459, p<0.0001, ηp^2^<=0.52; EE, right panel: phase × approach response: F_1,23_=3.8233, p>0.05] **(F).** EE influenced behavioral manifestations of anxiety on the elevated plus maze (EPM) **(G)** such that, despite spending a similar percentage of time in the open arms of an EPM [main effect of group: F_1,43_=1.472, p>0.05], EE rats made more head dippings while on the open arms than did SE rats [main effect of group: F_1,43_=3.8447, p<0.05, ηp^2^<=0.08]. *: p≤0.05.

We then tested whether the influence of housing conditions on these behavioral traits mediated the well-established effect of EE on the tendency to self-administer stimulant drugs (19–21). As previously described (21, 37), rats living under EE conditions were significantly less likely to acquire cocaine self-administration (SA) (**Fig. 2A**) than SE rats. However, the data also revealed that this EE effect was driven by a further decrease in the propensity to acquire cocaine SA in non-vulnerable populations, i.e. rats that have been shown either to have a low tendency to self-administer cocaine, characterized by a low locomotor response to novelty (low responders, LR) (7, 11) or a low saccharine preference (LSP) (38), or rats with low novelty (LNP) that are resilient to the transition to compulsive cocaine self-administration (12)(**Fig. 2B, 2C** and **Fig. S4-5**).

**Figure 2.**
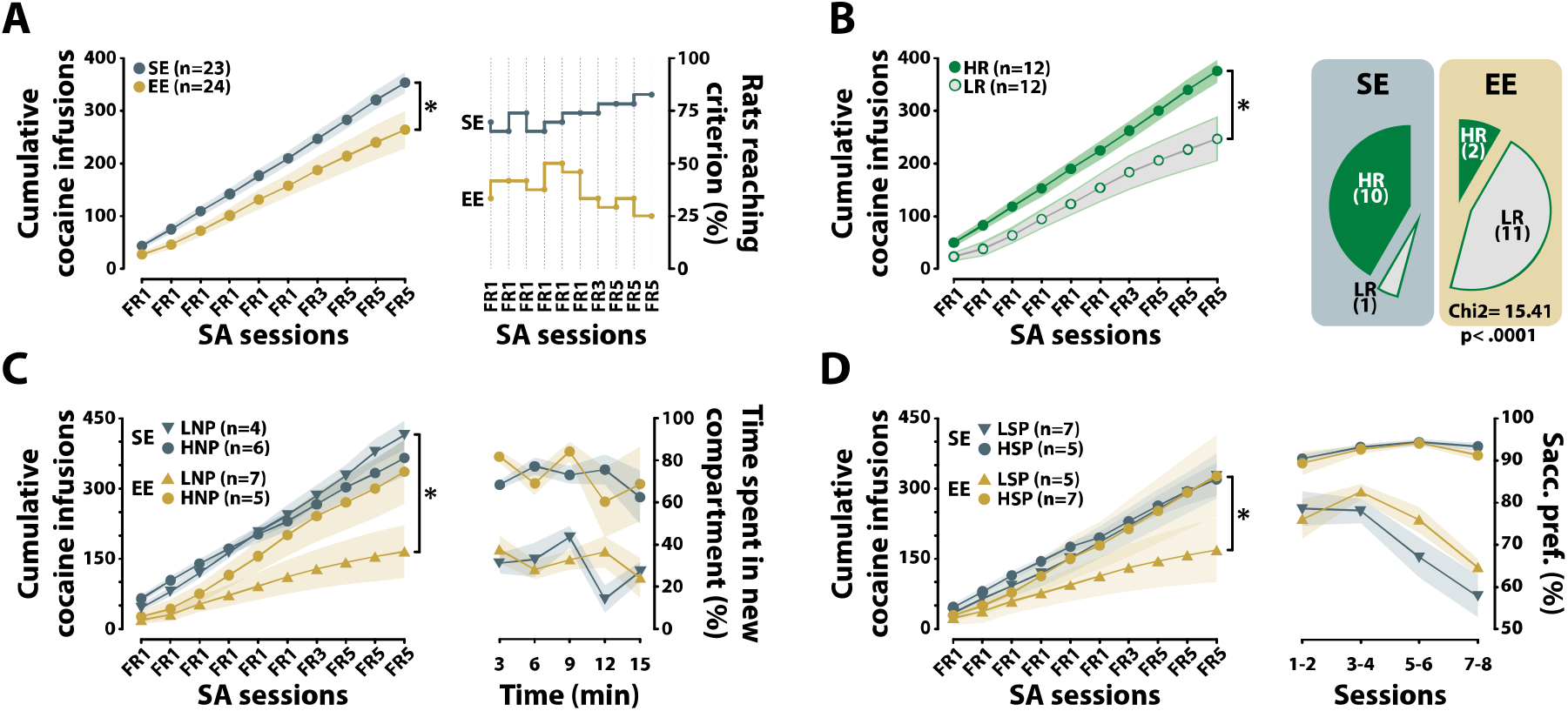
Environmental enrichment decreases the propensity to acquire cocaine self-administration in addiction-resilient rats. EE rats showed a lower rate of acquisition of cocaine self-administration (SA) than SE rats [main effect of group: F_1,46_=4.8168, p<0.05, ηp^2^<=0.10; session: F_9,405_=200.63, p<0.0001, ηp^2^<=0.82 and group x session interaction: F_9,405_=3.0058, p<0.01, ηp^2^<=0.06] (**A**, left panel). This effect was not simply driven by a small number of the EE population that failed to acquire SA. In fact, more than 50% of rats in the EE group stayed below the SA acquisition criterion (median number of cocaine infusions set at each session) (**A**, right panel). The high responder (HR) phenotype, which was almost non-existent in the EE population, violating the expected distribution typically observed in SE rats [Chi^2^=15.41, p<0.0001] (**B**, right panel), was the behavioral trait that best recapitulated the differential tendency to self-administer cocaine shown by EE rats [main effect of group: F_1,22_=7.1607, p<0.05, ηp^2^<=0.25; session: F_9,198_=138.78, p<0.0001, ηp^2^<=0.86 and group × session interaction: F_9,198_=3.7631, p<0.001, ηp^2^<=0.15] (**B**, left panel). EE also decreased the propensity of addiction-resilient, LNP rats, and rats with low reward sensitivity, LSP rats, to acquire cocaine SA [environment × phenotype × session interaction: F_9,162_=5.2896, p<0.0001, ηp^2^<=0.23 and F_9,180_=2.1008, p<0.05, ηp^2^<=0.10, respectively]. Thus, LNP and LSP rats from the EE population received far fewer cocaine infusions than their SE counterparts [planned comparison: p<0.01 and p<0.05 respectively] (**C** and **D**, left panels), even though the qualitative nature of these traits was not influenced by housing condition [environment × phenotype interaction: F_1,18_<1; environment × phenotype × time interaction: F_4,72_=1.9406, p>0.05, **C**, right panel and environment × phenotype interaction: F_1,20_=1.7919, p>0.05; environment × phenotype × time interaction: F_3,60_<1, **D**, right panel]. *: p≤0.05.

The increased weight of the addiction-like behavioral proneness conveyed by sweetness and novelty preference to the multidimensional behavioural profile shaped by EE, at the expense of locomotor reactivity to novelty which confers resistance to addiction-like behavior (7, 12) (**Fig. 2C**), suggests that while decreasing the tendency to self-administer cocaine, EE may make the switch from controlled to compulsive drug use more likely in vulnerable individuals.

We therefore investigated, in a large heterogenous cohort, the influence of housing conditions on the individual tendency to develop addiction-like behavior following prolonged exposure to cocaine SA (8, 12, 39).

### Environmental enrichment promotes the development of cocaine addiction-like behavior

Using new methodologies that enable the maintenance of i.v. catheter patency for several months in group-housed rats, we trained a new cohort of 48 rats (EE vs SE n=24 each) to self-administer cocaine for ~50 days before they were tested for their addiction-like behavior according to the stimulant addiction-relevant 3 criteria model (7, 8, 12, 39–41). This multidimensional model is based on the operationalization of the main behavioral characteristics of addiction in the DSM (9), namely high motivation for the drug, the persistence of drug taking despite adverse consequences and an inability to refrain from seeking the drug when signaled as unavailable (see (7, 39) and **Fig. S6**).

In order to determine the influence of housing conditions on the vulnerability to develop addiction-like behavior rats from both housing conditions were considered to form a large, heterogenous, cohort, and the individual tendency eventually to display each of the three addiction-like criteria was measured in that population irrespective of the housing conditions.

Regardless of their housing conditions, rats were identified as having 0, 1, 2 or 3 addiction-like behavioral criteria (**Fig. 3A and S6**, see **SOM**). As previously described (7, 39), the overall population was linearly distributed alongside an addiction severity index (8) in which only those rats showing the three addiction criteria (3-criteria or 3crit) had a score that was beyond the standard deviation of the cohort, while 0crit, addiction-resilient rats had a negative addiction severity score. As previously described (7, 12, 42, 43), the linear addiction severity index provides a criterion-dependent difference in the manifestation of each of the addiction-like behaviors so that 3crit addiction-vulnerable rats had scores significantly higher than those of 0crit addiction-resilient rats in each of the addiction-like behaviors (**Fig. S6A**).

**Figure 3.**
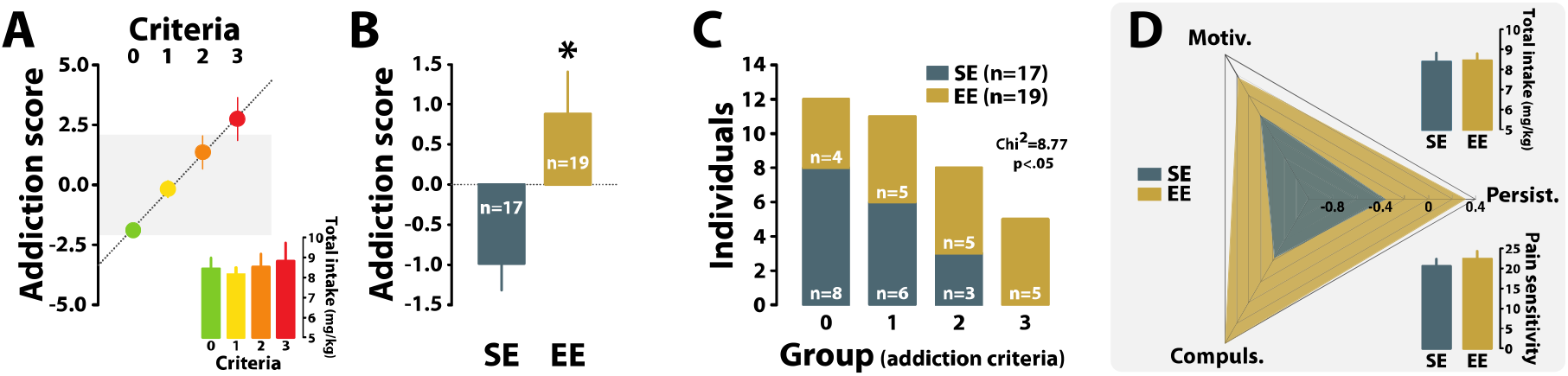
A negative contrast between housing conditions (EE or SE) and drug taking setting promotes the development of cocaine addiction-like behavior. After 47 days of cocaine SA under a FR-5 schedule of reinforcement, rats from a heterogenous cohort comprising similar numbers of individuals housed in SE or EE were tested for the three behavioral criteria of addiction-like behavior: high motivation for the drug measured under progressive ratio; the maladaptive engagement in drug seeking, measured as the inability to refrain from responding when the drug is signaled as unavailable; the persistence of drug seeking despite negative consequences, measured as resistance to punishment. Rats were stratified on the number of criteria they displayed, i.e. 0, 1, 2 or 3 criteria (**A**). The 3crit group (n=5, 14% of the entire population) was the only one having an average addiction score that was outside the standard deviation of the overall population (grey bar). In contrast, the 0crit group (n=12, 33% of the overall population) was the only one displaying highly negative scores confirming their resilient phenotype [main effect of crit: F_3,32_=17.955, p<0.0001, ηp^2^<=0.63] **(A)**. The difference between these groups was not attributable to a differential exposure to cocaine throughout their SA history [main effect of crit: F_3,32_<1]. However, retrospectively factoring housing conditions revealed that EE rats had significantly higher addiction scores than SE rats, the latter actually having negative scores [main effect of environment: F_1,34_=8.8255, p<0.01, ηp^2^<=0.21] **(B)**. All of the 3crit rats and the majority of 2crit rats were from the EE population, in clear contrast to the predominance of SE rats in the resilient, 0crit, population **(C)**. The higher addiction scores of EE rats as compared to SE rats were attributable to each of the addiction-like behaviours [main effect of environment: F_1,34_=8.82, p<0.01, ηp2=0.21, behavior: F_1,2_<1 and environment x behavior interaction: F_1,68_<1], even if more pronounced differences were observed for compulsivity and persistence of drug seeking than for the motivation for the drug. The facilitation of the transition to addiction by EE was not attributable to a differential exposure to cocaine or to a differential sensitivity to pain [main effect of environment: Fs_1,34_<1] **(D)**. *: p≤0.05.

The individual vulnerability to develop addiction-like behavior, seen in 13.8% of the population, was significantly influenced by housing conditions. A retrospective analysis revealed that EE rats had much higher addiction scores than SE rats (**Fig. 3B** and **S6B**) and that the entire 3crit population was comprised exclusively of EE rats (**Fig. 3C**). The influence of housing conditions on addiction vulnerability was primarily driven by a greater tendency to show compulsivity and the persistence of drug seeking, and less so by an increased motivation for the drug (**Fig. 3D** and **S6B**). Importantly it was not due to any overall differences in drug intake (**Fig. 3D** and **S6C)** nor to any differences in pain sensitivity (44) (**Fig. 3D**).

This dimensional approach provides clear evidence that while decreasing the tendency to initially engage in drug use EE instead increases the likelihood of developing addiction-like behavior in vulnerable individuals.

We further investigated whether the facilitatory effect of EE on the development of compulsive cocaine intake also extended to alcohol. In a third experiment, the tendency of rats housed in EE vs SE (n = 12 each) to develop quinine-resistant (45–48), compulsive, alcohol intake was assessed after several months of intermittent access to alcohol in a two bottle-choice procedure (49, 50).

EE rats did not differ from SE rats in their alcohol intake over 47 daily sessions of intermittent or unpredictable access to a choice between alcohol and water (**Fig. S7A**). However, as compared to SE rats, EE rats showed an increased tendency to relapse to alcohol drinking following several weeks of imposed abstinence (**Fig. 4A**) and did so compulsively in that they showed persistence of drinking at relapse despite adulteration by the bitter tastant, quinine (47) (**Fig. 4B** and **S7**). Ultimately, EE rats were shown to be more vulnerable than SE rats in their development of two key behavioral features of alcohol use disorder (2, 51).

**Figure 4.**
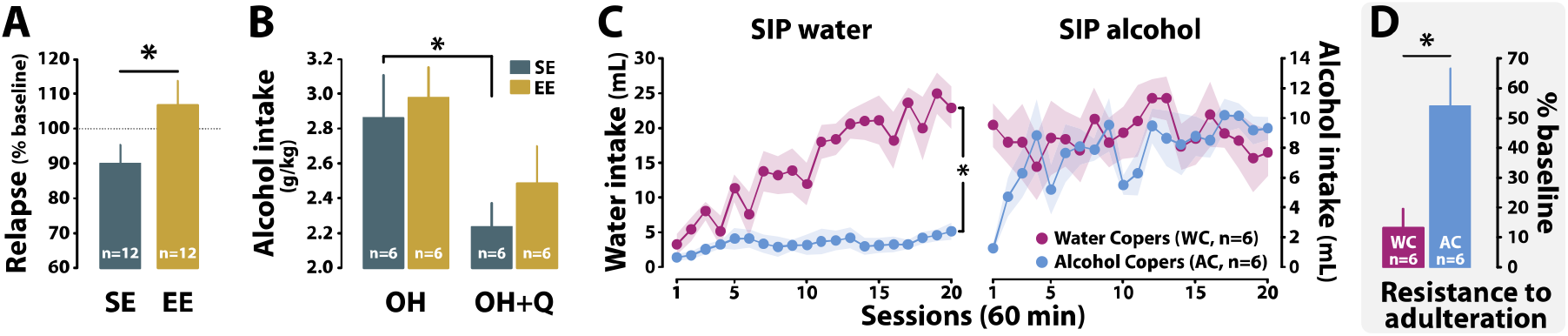
Alcohol drinking in a negative experiential state induced either by negative environmental contrast or by the aversive nature of the drug setting promotes the development of compulsivity. After 20 days of abstinence following a 3-month history of exposure to alcohol in an intermittent two-bottle choice procedure, EE rats were more prone to relapse to alcohol drinking than SE rats [Mann-Whitney: U=37.00, p<0.05] **(A)**. EE rats were also more prone to persist in drinking alcohol despite adulteration with quinine, thereby displaying compulsive drinking behavior [Kruskal-Wallis EE: H_1,12_=2.08, p>0.05; SE: H_1,12_=4.33, p<0.05] **(B)**. In another cohort of outbred SE rats, introduction of the opportunity to drink alcohol as a means of coping with distress in a schedule-induced polydipsia procedure resulted in a specific subpopulation of individuals (Alcohol Copers, AC) that developed adjunctive alcohol drinking behavior [main effect of group: F_1,10_=1.2771, p>0.05; session: F_19,190_=1.2348, p>0.05; group × session interaction: F_19,190_=1.4429, p>0.05; **C**, right panel] that they had failed to acquire when water was available, in marked contrast to rats that had acquired high levels of water intake (Water Copers, WC) [main effect of group: F_1,10_=33.619, p<0.001, ηp^2^<= 0.77; session: F_19,190_=9.8985, p<0.0001, ηp^2^<=0.50 and group × session interaction: F_19,190_=7.0845, p<0.0001, ηp^2^<= 0.41; **C**, left panel]. Although both WC rats and AC rats consumed the same amount of alcohol overall [main effect of group: F_1,10_=1.2771, p>0.05], only the latter, i.e. those that had acquired a coping strategy by drinking alcohol, subsequently displayed compulsive alcohol drinking, being resistant to adulteration of alcohol with quinine [main effect of group: F_1,10_=8.3820, p<0.05, ηp^2^<=0.46] **(D)**. *: p≤0.05.

Together, these results reveal a bidirectional effect of housing conditions on the initial tendency to self-administer drugs and on the vulnerability to compulsively take drugs (2). The only difference between EE and SE rats in these parametrically controlled experiments was the enrichment of living context of the former and the resulting differential negative contrast with the drug taking context (**Fig. S8)** shown previously to influence the relative preference for cocaine or heroin (52). Thus, while these results may at first glance appear counter-intuitive, or at variance with the literature on the effects of environmental enrichment on drug related behaviors based on the comparison of EE and SI rats (see 37 for further discussion), they are highly consistent with previous evidence that EE rats are more sensitive to the motivational effects of cocaine than SE rats (22) and highlight the importance of the experiential factors inherent in the contrast between a rat’s living conditions and the drug-taking setting.

In contrast to the well-known ‘Rat Park’ experiment, in this study EE rats were provided access to cocaine or alcohol in a relatively impoverished drug setting, rather than in the enriched housing environment itself (53). Additionally, the “Rat Park” experiment did not investigate individual differences in addiction-relevant behaviors, such as compulsive drug intake, that emerge only after a prolonged period of training. In the present study EE rats, unlike SE rats, first encountered the drug in a state of relative social, sensory and cognitive deprivation because of the marked contrast between their enriched housing conditions and the drug SA context the features of which were similar to those of the SE (**Fig. S8**). This negative environmental contrast likely biases individuals to self-administer drug in the experiential setting to ameliorate an internal deficit/distress state (5, 54). In humans, this negative motivational state is frequently associated with the initiation of drug use and increases the vulnerability to addiction (4, 55) by moderating or exacerbating the behavioral traits that here we show are influenced by EE, namely sensation seeking, boredom susceptibility/novelty preference and anxiety (55–57).

According to this framework, rather than solely an animal’s housing conditions, it is the experiential factors in the drug setting, such as those related to high boredom susceptibility in humans or novelty preference in rats (12, 58, 59), that influence the vulnerability subsequently to compulsively take drugs. We therefore extended the study to causally test the hypothesis that the acquisition of alcohol drinking as a coping, stress-reducing, strategy is sufficient to exacerbate the vulnerability to switch to compulsive alcohol intake in rats living under similar housing conditions.

The acquisition of adjunctive behaviors, such as schedule-induced polydipsia (SIP), has been shown across species to result in a profound decrease in the levels of stress-related hormones that are increased by intermittent food delivery (60, 61), thereby providing a valuable tool to investigate the influence of alcohol drinking as a coping strategy on the subsequent vulnerability to develop compulsive alcohol intake. In SIP procedures, most rats readily acquire an adjunctive, anxiolytic, water drinking response (62–64) (**Fig. 4C**), as a way of coping with distress. However, some rats never develop this adjunctive behavior (62–64). We hypothesized that these individuals might instead develop adjunctive alcohol, not water, drinking as a coping response. We further predicted that rats having selectively acquired adjunctive alcohol drinking would also go on to develop compulsive (quinine-resistant) alcohol intake (see **SOM**)..

In a fourth longitudinal experiment, alcohol was provided to a cohort of 48 SE rats (65) that had been trained to cope with the distress of SIP by adjunctively drinking water (61, 66, 67). A sub-population of these rats also maintained their established coping (adjunctive drinking) strategy when alcohol was introduced in place of water (Water Copers, WC). However, as predicted, a distinct sub-population of rats never acquired adjunctive water drinking and only developed a coping strategy by drinking alcohol (Alcohol Copers, AC) (**Fig. 4C**).

AC rats, which did not differ from WC rats in terms of total alcohol intake over the course of the 20 days of exposure (**Fig. S9** and **SOM**), also persisted in drinking alcohol when adulterated with quinine (**Fig. 4D** and **S9**). This differential vulnerability to develop compulsive alcohol drinking was not due to any differences in blood alcohol levels since they were similar in both groups (**Fig. S9**) and it was therefore specific to the experiential nature of the SIP context. Only rats that had acquired the adjunctive, anxiolytic behavior of drinking alcohol went on to drink alcohol compulsively.

Together these results show that a negative experiential context, whether inherent to the drug taking environment or triggered by a negative affective contrast from housing/living conditions at the onset of alcohol use, may be an important gateway to the development of compulsive drinking in individuals unable to learn to cope with negative states by alternative means (**Fig. 4D** and **S9**).

## Discussion

The present findings provide clear evidence for the role of environmental and experiential factors in shaping an individual’s vulnerability to make the transition from controlled to compulsive drug taking.

In replicating the well documented effect of EE on locomotor reactivity to novelty (68), anxiety (69), sign tracking (33) and stimulant SA (19) this study further showed that the decrease in cocaine intake shown by EE rats is associated with the loss of the HR trait, a factor of resilience to addiction (7, 12, 30). The influence of EE on the multidimensional behavioral profile of the individuals observed here is consistent with previous evidence that the environment shapes, but does not determine, individual differences in drug-related behavior within a large cohort of individuals (70). Furthermore, the effects observed here cannot be attributable to the influence of social isolation, so often used as a control condition in earlier experiments (37).

In contrast to these previous studies and experiment 1, rats in experiment 2 were trained to self-administer cocaine intravenously over prolonged periods of time (over two months). The new self-i.v. administration methodology we developed for this study enabled the investigation of the influence of different housing conditions over the peri-adolescence period and throughout the long history of drug self-administration on an individual’s vulnerability to develop compulsive drug taking.

While confirming their lower tendency to acquire drug self-administration, EE rats were shown to be more, not less, likely than SE rats to develop compulsive (punishment- (7, 8, 47) or adulteration-resistant (46, 47)) drug intake, that was independent of any differences in total drug intake. The dissociation between overall exposure to the drug and the transition to addiction-like behavior confirms our previous demonstration that addiction-like behavior for cocaine develops only in a subset of individuals having different patterns, but not different levels, of drug intake (8, 12, 39). Additionally, the escalation of cocaine intake, which has been shown not to differ between EE and SE, as opposed to SI rats (17), has been shown to be a consequence of, and not a causal contributor to, the development of compulsive cocaine SA (8, 71). Similarly, the preference to drink alcohol that characterizes P rats and their overall history of alcohol intake did not predict the subsequent individual differences in the tendency to compulsively seek alcohol (72, 73).

The parsimonious interpretation of these observations involving drugs from two distinct classes is that the vulnerability to develop compulsive, punishment-resistant, cocaine and alcohol intake – is exacerbated by the negative contrast in EE rats between their enriched housing conditions and the minimal, or impoverished, environment of the drug taking context. Despite previous evidence that rats living in large cages display a central response to stress not shown by rats living in small environments (74), the nature of the cognitive, sensory, and emotional deprivation that individuals experience when tested for several hours in relatively impoverished conditions outside their enriched cages has not hitherto been considered. It is nevertheless revealed here to have the marked effect of exacerbating an individual’s vulnerability to develop compulsive drug taking.

This observation resonates with evidence that, irrespective of their socioeconomic status, the affective state in which individuals engage in drug use is an important determinant of their vulnerability to addiction (4). Drug use as an emotion regulation, coping, strategy, has long been suggested to be a key determinant of the transition to addiction, highlighting the importance of experiential, rather than pharmacological, factors in the vulnerability to switch from controlled to compulsive drug use (5, 54, 75, 76).

We therefore tested the hypothesis that it is the experiential factors associated with the acquisition of drug use in the drug setting that are important determinants of the development of compulsive drug intake. To do so, we developed a novel procedure that draws on the marked individual variability in developing anxiolytic, adjunctive drinking behavior in a schedule-induced polydipsia procedure (77). Rats that develop adjunctive drinking in a SIP procedure are also better able to establish avoidance strategies than those that do not (78) thereby revealing a generally greater ability adaptively to cope with distress. We show here that these rats continue to drink alcohol adjunctively when water is replaced by alcohol. We and others have also established that some rats do not readily engage in adjunctive water drinking (62, 64) as a stress-reducing response. We therefore hypothesised that rats that do not develop adjunctive water drinking when exposed to the stress-inducing intermittent food delivery of a SIP procedure would instead be able to develop an adjunctive response by drinking alcohol. If this proved to be the case, we predicted that alcohol-SIP rats would also show compulsive, quinine-resistant, alcohol drinking. We further predicted that those rats that had developed an adjunctive response with drinking water and continued that behavior when water was replaced by alcohol would suppress their alcohol drinking when adulterated with quinine, i.e. would not drink compulsively.

The results of experiment 4 confirmed our predictions in that water-copers, rats characterized by their ability to cope with distress, continued to express an adjunctive response when water was replaced by alcohol, but they did not show resistance to adulteration with quinine. In contrast alcohol-coper rats that had selectively acquired an alcohol, but not water, adjunctive drinking response, also eventually went on to be resistant to quinine and persist in drinking. This compulsive drinking was specific to the adjunctive context, as they no longer differed from the control group when tested for their resistance to adulteration in their home cage.

The development of high levels of drinking shown by alcohol-copers cannot be attributed to delayed development of SIP as we have previously shown that low drinkers for water maintain a very low level of water intake over an even longer period of time (79). These results therefore show that experiential factors associated with the acquisition of alcohol use as an adjunctive response, and not the level of intoxication or alcohol exposure, can exacerbate the individual tendency to develop compulsive alcohol drinking.

## Conclusions

Together, the results of these experiments show that considering the drug setting or the living environment alone when trying to understand the nature of the gene × drug × environment interactions that predispose vulnerable individuals to become addicted to a drug does not fully capture the importance of the experiential factors operating during initial exposure to the drug. These factors themselves depend on the interaction between housing/living conditions and the drug taking setting. While providing further evidence for a previously under-estimated role of non-pharmacological factors, highlighted by Ahmed and colleagues (80), in the vulnerability to progress from controlled to compulsive drug taking, the present results further suggest that initiating drug use through negative reinforcement-based self-medication (5) (from the “dark side” (81)), is an important determinant of the development of addiction in vulnerable individuals.

## Materials and Methods

This study comprised 4 independent experiments involving 192 male Sprague Dawley (SD) rats (for a detail of the methods, see **SOM**). Experiment 1 was conducted to test the hypothesis that the effect of housing conditions (EE vs SE) on the propensity to acquire cocaine self-administration (SA) was mediated by their influence on the behavioral traits of vulnerability or resilience to addiction and the associated propensity to acquire cocaine SA. Having reproduced the well-established observation that EE decreased the propensity to acquire cocaine SA, we further demonstrated that this effect was mostly driven by resilient individuals, e.g. rats previously shown to be less likely to develop compulsive cocaine self-administration such as high responder to novelty (HR) or low novelty-preference rats. We were then able to investigate the influence of EE on the transition from controlled to compulsive drug use in vulnerable individuals. Thus, we tested the influence of environmental conditions on the development of addiction-like behavior by exposing rats to ~ 50 days of cocaine SA and assessing their score in the 3-criteria model of cocaine addiction (42).

Because this experiment demonstrated that EE promotes the development of addiction-like behavior, we tested the hypothesis that this effect is not restricted to cocaine but also to alcohol. Having demonstrated that this facilitatory effect of EE on the development of compulsive drug intake was also seen in rats drinking alcohol. Thus, we demonstrated that this facilitatory effect of EE on the development of compulsive drug intake was also seen in rats drinking alcohol. We hypothesized that the facilitation of the transition to addiction by EE was dependent on the negative contrast between EE housing conditions and the relatively impoverished drug taking setting, thereby biasing the acquisition of drug taking as ‘self-medication’ of a deprived state. The final experiment was conducted to test the hypothesis that if individuals were to initiate alcohol use as a coping strategy in response to a distress independent of a contrast between housing and testing conditions, as measured under schedule-induced polydipsia conditions (82), they should also be more vulnerable to develop compulsive alcohol drinking than individuals that had not have needed to drink alcohol to learn to cope with distress.

### Experiment 1

Following a week of habituation to the vivarium, forty-eight 6-7 week-old male SD rats were housed in EE (n=24) or SE (n=24) conditions (**see SOM**) until the end of the experiment. Subsequently, their sensitivity to a natural reward (83) was assessed in a two-bottle choice procedure with water and saccharine (**Figure S1**). Rats were then screened for their anxiety-related behavior and their locomotor reactivity. The individual variability in ascribing incentive value to a CS and novelty preference was measured in an autoshaping (sign-tracking) (30, 84) and a novelty-induced preference task (12).

At day 96, rats underwent surgical implantation of an indwelling catheter in the right jugular vein for SA training. Five days later individual differences in the acquisition of cocaine SA were assessed, starting with six 3.5-hour daily sessions under a fixed ratio (FR) 1 schedule of reinforcement (FR1) followed by one under FR3 session and three under FR5.

A normalized score for each individual for each trait (% time spent in the open arm (OA), saccharine preference (%), time spent in novel compartment (%), total distance travelled in the open field (OF) and contacts made with the CS-associated lever) was calculated by subtracting the mean of the population to each individual value and dividing the result by the standard deviation of the population (z-score) as previously described (42). The z-score of each rat was used within a multi-dimensional model (represented as radar plots) within which the contribution of each behavioral trait to the multidimensional behavioral characterization of each individual (referred to as rat multidimensional behavioural profile) was calculated as the percentage of the normalized score of all the behavioral traits.

These multidimensional behavioural profiles were subjected to a k-means cluster analysis (72, 85) performed on the value (percentage) of each of the 5 behavioral traits with the following parameters: maximization of distances between clusters, maximum 20 iterations, maximum of 3 clusters.

### Experiment 2

Following a week of habituation to the vivarium, forty-eight male SD rats were housed in EE (n=24) or SE (n=24) conditions until the end of the experiment (**Figure S1**). Five days following the implantation of an indwelling catheter into their right jugular vein, rats were trained to self-administer cocaine for 5 sessions under an FR1 schedule of reinforcement followed by 2 sessions of FR3 and 40 sessions under a FR5 schedule of reinforcement.

The individual vulnerability to develop addiction-like behavior was assessed as previously described (7, 12, 40, 42, 43) by (i) the inability to refrain from responding despite the drug being signalled as unavailable, during so-called no-drug periods, (ii) a high motivation for cocaine, assessed over a single session under a progressive-ratio schedule of reinforcement (40) and (iii) the persistence of responding for cocaine despite punishment, as measured in a single 40 min long session during which instrumental responses were punished response-contingently (40).

Animals were scored for each addiction-like behavior independently. If the score of the individual was in the 33% highest percentile of the distribution, that individual was considered positive for that criterion. Animals were then separated in four groups depending on the number of positive criteria met (from 0 to 3). An addiction score was calculated for each animal as the sum of the normalized scores (Z-scores) for each of the three addiction-like criteria (12, 42).

### Experiment 3

Following a week of habituation to the vivarium, twenty-four male SD rats were housed in EE (n=12) or SE (n=12) conditions until the end of the experiment (**Figure S1**). Rats were first exposed to 21 two-bottle choice sessions under intermittent access. Rats were subsequently exposed to 20 two-bottle choice sessions under unpredictable to promote binge-like drinking behavior followed by 6 sessions under intermittent access. Access to alcohol was then withdrawn for 20 days during which rats stayed in their home cage with access only to water. Rats were subsequently challenged with a relapse test which consisted of a first 4-hour two-bottle choice session followed by a 24-hour two-bottle choice session. The individual propensity to relapse was calculated as the ratio between alcohol intake at relapse and alcohol intake in the last 4-hour two-bottle choice session prior to abstinence.

EE and SE rats were then divided into two subpopulations (n=6 each) displaying the same drinking behavior at every stage of the experiment (training pre-abstinence and during relapse). One subpopulation of EE and SE rats was exposed to two sessions during which alcohol was adulterated with quinine (0.1 g/L) (86) while the other subpopulation was exposed to non-adulterated alcohol. This between-subject approach was carried-out to control for any potential confounding influence of a habituation to the bitter taste of quinine (latent inhibition) from which a within-subject design may have suffered. Finally, to control for the specificity of the effect of quinine on alcohol intake, rats underwent two 4-hour water adulteration sessions during which they had access to a single bottle of water supplemented with quinine (0.1 g/L).

### Experiment 4

Following a week of habituation to the colony, forty-eight male SD rats were all housed in a SE (**Figure S1**). Rats were food restricted to 80% free-feeding body weight (62, 64) for a week before being trained in a schedule-induced polydipsia (SIP) procedure with water for twenty 1-hour daily sessions, as previously described (62, 64). Then, water was substituted with 10% alcohol and rats were trained in twenty 1-hour daily SIP sessions with alcohol. The day following the last SIP session with alcohol, the individual tendency to persist in drinking alcohol despite adulteration with quinine (0.1 g/L) was assessed.

Rats then underwent five baseline SIP with alcohol sessions over which they recovered their pre-adulteration alcohol intake level. Blood samples were collected immediately after the last session to measure their BAL.

In order to test whether persistence of drinking alcohol despite adulteration by rats trained under the SIP procedure was dependent on the SIP context in which rats initiated their alcohol use, another cohort (n=24) was housed under similar conditions and was then given a two-bottle choice procedure under intermittent access for twelve 24-hour sessions, followed by twelve 1-hour sessions. Rats were subsequently given eight additional 24-hour two-bottle choice sessions under intermittent access followed by a single 1-hour session during in which their individual tendencies to maintain alcohol drinking despite adulteration was measured.

The individual propensity to persist in drinking alcohol despite adulteration was calculated as the ratio between “alcohol + quinine” intake over the average alcohol intake of the first hour during the last three two-bottle choice procedure sessions under intermittent access.

### Data and statistical analyses

Data are presented as number of individuals, mean (for some radar plots) or mean +/− standard error of mean (SEM). Behavioral data were subjected to repeated measure, one-, two-factorial analyses of variance (ANOVAs) or Kruskal-Wallis or Mann-Whitney U wherever appropriate. The experimental groups were used as between-subject factor and, depending on the analysis, the arm (open/closed arms) and the time (number of sessions or session blocks) were used as within-subject factors. Upon confirmation of significant main effects differences among individual means were analyzed using the Newman-Keuls post-hoc test and planned comparisons wherever appropriate. a was set at 0.05 and the effect size was reported by the partial eta-squared value (ηp^2^) (see **SOM** for more details).

## Acknowledgments

This work was supported by a Programme Grant from the Medical research Council to BJE and DB (MR/N02530X/1), an INSERM AVENIR grant and a research grant from the Leverhulme Trust to DB (RPG‐2016‐117) as well as a Leverhulme Trust Early Career Fellowship to LMP (ECF-2018-713).

