## Supplemental material for "Environment-dependent behavioral traits and experiential factors shape addiction vulnerability"

**This PDF file includes:**

Supplementary methods

Supplementary text

Figures S1 to S9

SI References

Supplementary Methods

### Subjects

One hundred and ninety-two male Sprague Dawley rats (*Charles River laboratories, France or UK*), 6/7 weeks old, weighing approximately 200g upon arrival were housed 2 per cage for a week of habituation in a standard environment (40.5x35.5x23cm ventilated cages). Rats had ad libitum access to food and water and were maintained under a reversed 12-hour light/dark cycle (light ON between 7.00pm and 7.00am). Rats were then food restricted (15-20g chow daily) and tested in the behavioral procedures detailed below (6-7 days/week). Procedures were conducted in accordance with the European Community Council Directives (86/609/EEC) and with the United Kingdom Animals (Scientific Procedures) Act 1986, amendment regulations 2012 following ethical review by the University of Cambridge Animal Welfare and Ethical Review Body (AWERB) PPL70/8072.

### Environmental enrichment

Following a week of habituation, rats (see “specific experimental timelines”) either remained housed 2 per cage in standard cages, referred to as standard environment (SE) throughout the manuscript or were housed in groups of 4 in an enriched environment (EE) consisting of a large cage (80x50x80cm, *Ferplast*) equipped with toys (tubes, blocks), a running wheel, platforms, houses and hammocks. The toys and the configuration of the different elements within each cage were changed weekly (1).

**Characterisation of behavioral traits**

Anxiety-related behavior, novelty-induced place preference and locomotor reactivity to novelty were measured using a video tracking system (*ViewPoint Behavior Technology®*, *Lyon, France*) on an Elevated Plus Maze (EPM) (2), four Novelty Place Preference (NPP) boxes (3) and four Open Fields (OP) (4), respectively. Individual differences in incentive salience attribution to conditioned stimuli were assessed in operant chambers in an AutoShaping task (5) and sensitivity to natural rewards was conducted in a two-bottle choice procedure, water vs saccharine (6).

#### *Anxiety-related behavior*

High anxiety trait measured on an EPM has been shown to predict the escalation of cocaine SA (2).

The EPM is elevated 50cm above the floor and comprised two open arms (45x10cm), two closed arms (45x10x45cm) and a central platform (10x10cm). The luminosity was measured in each portion of the EPM and the light adjusted to reach an intensity of 40 Lux in the central platform, 50 Lux in the open arms and 30 Lux in the closed arms (2). The tests were conducted as previously described (2, 4) during the dark phase of the dark/light cycle between 8am and 11am.

At the beginning of a test session, rats were placed on the central platform with their head pointing towards an open arm (always the same arm for all subjects). They were then allowed to explore the maze for 5 minutes while their movements were monitored. In addition to entries into, and time spent in, each arm, exploratory behaviors, i.e. head dipping in the distal part (last 10cm) of open arms, were recorded as a measure of anxiety-related behavior (7).

An anxiety score, used in the multidimensional behavioural model, was calculated for each subject as a percentage of time spent in the open arms over the time spent in all arms. Individuals belonging to the upper and lower quartiles (25%) of the population were considered Low Anxious (LA) and High Anxious (HA), respectively (2).

#### *Novelty-induced place preference*

Novelty-induced preference has been shown to identify rats that are vulnerable (high novelty preference, HNP) or resistant (low novelty preference; NLP) to developing compulsive cocaine SA (3, 8).

The novelty-induced place preference (NPP) test was performed in four conditioned place preference boxes consisting of one narrow central corridor (10 x 50 x 50cm) surrounded by two large compartments (20 x 50 x 50cm), as previously described (3). Two guillotine doors located at both extremities of the central corridor gave access to the two large compartments protected by Plexiglas lids in order to decrease the luminosity to 0-0.2 Lux as compared to 10 Lux in the central corridor. The configuration of the boxes was adjustable such that the walls and floors could either be black or white and their texture smooth or rough. The different configurations were randomized between individuals (3).

The single test session for each rat, conducted during the dark phase of the dark/light cycle (7.00am-7.00pm), consisted of three successive steps as previously described (3). The animal was first placed in the central corridor with the two doors closed for a short habituation period of 5 min after which it was placed in one of the two compartments, considered the “familiar compartment” (randomized between individuals) for 25 min. In the final step, the animal was placed in the central corridor 5 seconds before the two doors were opened, giving free access to all the compartments of the box for 15 min.

The time spent in each of the 2 compartments (familiar and novel) was recorded and a score of NPP was calculated for each subject as the percentage of time spent in the novel compartment over the time spent in the other two compartments. Individuals in the upper and lower quartiles (25%) of the population were considered as HNP and LNP rats, respectively (3).

#### *Locomotor reactivity to novelty*

High locomotor reactivity to novelty has been shown to reflect individual differences in the activation of the stress system in response to a new inescapable environment (9) and has long been suggested to represent a subdimension of the sensation seeking trait (10). Additionally, the High Responder phenotype (HR) has been shown to be a behavioral marker of the increased propensity to acquired drug SA as well as a marker of resistance to addiction-like behavior to cocaine (3, 11).

The locomotor reactivity to novelty test was conducted as previously described (4) in four open fields (50 x 50 x 50cm) during the light phase of the dark/light cycle (7.00pm-7.00am) under a light intensity similar to that of the holding rooms (~ 550 Lux at the centre of the open field). Rats were placed in the open fields for two hours and their locomotor activity recorded throughout. Individuals whose total distance travelled was in the upper and lower quartiles (25%) of the population were considered High Responders (HR) and Low Responders (LR) rats, respectively.

#### *Pavlovian conditioned approach: Sign tracking and Goal Tracking*

Two days prior to the beginning of autoshaping training for food (i.e. 45 mg food pellets, *Bio-Serv, USA*), food-restricted rats received 20 pellets in their home cage in order to minimise any food neophobia.

The experimental sessions were conducted for 6 consecutive days in 12 operant chambers (31.8 cm x 25.4 cm x 26.7 cm each, *Med associates, St. Albans, USA*) located within a ventilated sound-attenuating cubicle. Front and back panels of the test chambers were aluminium, while right and left walls and the roof were transparent acrylic plastic with stainless grid floor. A pellet dispenser was installed behind the front wall of each chamber, supplying food pellets to a food magazine located 2 cm above the grid floor. Each test chamber was illuminated by one 3-watt light bulb (house light) during the experimental session. The first day of training was a habituation session in which rats were placed in the operant chamber (house light and fan turned on) for 30min. The second day of training was a single session of “magazine training” for the rats to learn where food pellets were delivered in the experimental chambers. In this session, 60 pellets were delivered according to a 30s-variable time (VT) schedule. Rats were then trained to associate the presentation of a compound conditioned stimulus (CS) comprising a cue-light and a lever, deemed active in that it enabled the measurement of Pavlovian approach and contacts with the compound CS. Responding on that lever had no programmed consequences to the subsequent delivery of a food pellet (unconditioned stimulus: US); i.e. there was no instrumental contingency.

CS-US presentations were delivered according to a VT a Variable time (VT) 90 seconds schedule (VT-90s) such that upon termination of each variable time, the Cue light was illuminated and the active lever was inserted for a 8 seconds period, after which the CS was turned off, the active lever was retracted, and one pellet was delivered in the food magazine. During the 8 seconds interval, contacts with the compound CS measured as active lever depressions, were recorded as sign-tracking events while head entries into the food magazine were recorded as goal-tracking events (5). Inactive lever presses (uncoupled from the CS) were also recorded as an indicator of general activity. Individuals within the upper and lower quartiles of the population stratified on their level of sign tracking (lever presses) over the last three sessions, were considered sign-trackers (ST) and goal trackers (GT), respectively.

#### *Saccharine preference*

The saccharine preference test was carried out over 12 daily sessions, as previously described (6). Prior to testing, rats were water deprived for 2h in order to avoid any influence on performance of individual differences in thirst. Rats were weighed immediately before the beginning of each session in order to be able to compute a fluid intake/body weight ratio. The two-bottle choice sessions were conducted every day, in cages similar to that used for SE housing with the exception that no bedding was present.

After four habituation sessions during which rats were given access to two bottles of water, they were then given the choice between one bottle containing water and one bottle containing 0.2% saccharine (*Sigma*) for eight 2-hour sessions. Bottles were weighed immediately before and after each session to measure the intake of both solutions. One hour following the beginning of each test session, bottles were swapped for the remaining 1-hour period to control for the development of a side preference bias. The ratio between saccharine intake over the total fluid intake was calculated, and individuals belonging to the upper and lower quartiles of the population stratified on saccharine preference over the sessions were considered as High saccharin Preference (HP) and Low saccharin Preference (LP) rats, respectively (6).

### Surgeries and drugs

Rats were anesthetised by intraperitoneal injection of a ketamine/xylazine mixture (*Ktamine1000, 90 mg/kg, Virbac, France; Rompun 2%, 6.7 mg/kg, Bayer, France*) and a silastic catheter (*CamCaths, UK*) was implanted into the right jugular vein as previously described (12). Rats were treated for one week with daily subcutaneous injection of an antibiotic (*Baytril, Bayer*) and catheters were flushed daily with 100μl of sterile physiological saline (0.9% sodium chloride) supplemented with heparin (50 IU/ml).

Cocaine hydrochloride (*Coopération Pharmaceutique Française*) was dissolved in sterile physiological saline (0.9% sodium chloride) at a final concentration of 2.5g/L and stored at 4°C.

### Cocaine self-administration

Cocaine self-administration (SA) was conducted as previously described (12) in operant chambers made of aluminium and transparent acrylic plastic with stainless grid floor (31.8 cm x 25.4 cm x 26.7 cm, *Med associates, St. Albans, USA*) located within a ventilated sound-attenuating cubicle. Each chamber contained two retractable levers (4cm wide), located on the front panel such that rats had access to both an active and an inactive lever, the location of which was randomised between individuals. Each test chamber was illuminated by one 3W light bulb (house light) during the experimental session and cue lights (2.5W) located above each lever. An additional cue light (central cue light, 2.5W) was located above and between the lever-associated cue lights.

Flexible Tygon tubing, protected by a metal spring anchored to a swivel secured on a pivoting arm, linked a perfusion pump located outside the chamber at one end to the external port of the catheter on the other side. The stainless steel grid floor was linked to a scrambler (*Med associates, St. Albans, USA*) for the delivery of mild electric foot-shocks as previously described (13). All the SA experiments were performed 6/7 days a week during the dark phase of the light/dark cycle.

#### *Intravenous cocaine self-administration training*

SA training was performed as previously described (3, 14-16). Chiefly, daily SA sessions comprised three 40 min periods during which the drug was signalled as available by a discriminative stimulus (house light ON) separated by two 15-min drug free periods (called no-drug periods) during which the house light was turned off. During the "no-drug" periods, lever presses were without scheduled consequences. During the "drug" periods, active lever presses under a Fixed Ratio (FR) schedule of reinforcement resulted in the illumination of the drug-paired cue light (CS) for 5s and the activation of the infusion pump. Inactive lever presses had no scheduled consequences. Each infusion (0.25mg/100µL/5.7s) was followed by a 40sec time-out period. Rats acquired SA over the first 5-6 days under a FR1 schedule of reinforcement after which the FR was incrementally increased to 3 and then 5 over three days (14).

#### *Progressive-ratio schedule*

During the progressive-ratio schedule of reinforcement, drug availability was signalled by the illumination of the house light. The ratio of responses per infusion was increased after each infusion according to the following progression (10, 20, 30, 45, 65, 85, 115, 145, 185, 225, 275, 325, 385, 445, 515, 585, 665, 745, 835, 925, 1025, 1125, 1235, 1345, 1465, 1585), as previously described (3, 14-16). The maximal number of responses made by a rat to obtain one infusion determined the last ratio completed, which is referred to as the breaking point. The session ceased after either 5 hours or when a period of 1 hour had elapsed since the previously earned infusion.

#### *Punishment sessions*

The propensity of rats to persist in responding for cocaine despite punishment was tested for the duration of the first drug period, i.e. 40 min, as previously described (3, 14-16). The house light signalled that the drug was availability. The schedule was the following: FR1 led to the illumination of the central cue light signalling the presence of the shock. On completion of an additional 3 lever presses, thereby reaching FR4, a mild electric shock was delivered (0.45 mA, 2 sec). On completion of a fifth lever press, thereby completing the ratio requirement of 5 (FR5), rats received an electric shock (0.45 mA, 2 sec) followed by a cocaine infusion associated with the presentation of the CS for 5s. Then the central cue light was turned off.

If, within a minute following the first lever press (FR1), rats did not complete an FR4 or, after having received the shock following the fourth lever press, did not complete the FR5 sequence, the central cue light turned off and the sequence was reinitiated (14).

**Pain sensitivity measurement**

To ensure that potential differences in compulsivity were not attributable to a differential sensitivity to pain, rats were tested on a hot plate prior to the punishment sessions. Six hours following the self-administration session, rats were placed on a hot plate (*Ugo Basile, Gemonio, Italy*) set-up to remain at a stable temperature of 52ºC (17). The time elapsed before the appearance of pain-associated behaviors, including paw-licking and jumping, was measured and considered an indication of pain threshold (17). Rats were then immediately removed from the hot plate and returned to their home cage.

### Two-bottle alcohol-water choice procedure

Rats and bottles were weighed prior to, and following, each session in order to compute a fluid intake/body weight ratio. The two-bottle choice sessions (alcohol versus water) were adapted from Simms et al. 2008 (18) and were conducted in cages similar to that of the standard environment (40.5 x 35.5 x 23cm ventilated cages).

#### *Intermittent access to alcohol*

To maximize alcohol consumption, rats were exposed to an intermittent two-bottle choice procedure adapted from Simms et al. (18). Rats had free access to two bottles containing either alcohol or water for twenty-four hours followed by twenty-four hours of abstinence (rats were returned to their home cages). The location of the alcohol bottle was changed over the course of each session to control for potential development of a side preference. The first four sessions consisted of habituation to alcohol during which the ethanol concentration was increased from 3% to 20% (3, 6, 10 and 20%). Rats then underwent a specific number of sessions (see specific experimental timelines) during which they had free access to water or 20% alcohol.

#### *Unpredictable access to alcohol*

In an attempt to elicit binge-like alcohol drinking, unpredictable access to alcohol was introduced in which the duration of each session was variable and unpredictable (from 30 minutes to 12 hours). The two-bottle choice sessions were separated by twenty-four hours of abstinence.

#### *Alcohol adulteration*

During the alcohol adulteration sessions, rats were given free access to a single bottle of 20% alcohol containing 0.1 g/L quinine (Sigma-Aldrich) (19) for four hours. In order to control for the specificity of the effect of quinine on alcohol intake, rats were also exposed to two 4-hour water adulteration sessions during which they had access to a single bottle of water containing quinine (0.1g/L).

### Schedule-induced polydipsia

The Schedule-induced polydipsia (SIP) procedure was carried-out as previously described (20, 21). Rats were trained daily in operant chambers made of aluminium and transparent acrylic plastic with a stainless-steel grid floor (24 cm x 25.4 cm x 26.7 cm, *Med associates, St. Albans, USA*) located within a ventilated sound-attenuating cubicle. Chambers were equipped with a house light (3-W), a food tray (magazine) installed at the center of the front wall, and a bottle from which a stainless-steel sipper tube delivered water into a receptacle placed in a magazine on the wall opposite the food magazine. Water or alcohol were freely available throughout the 42 1hour sessions.

#### *Habituation / baseline water intake*

During the habituation session, rats were given access to water and 60 food pellets (45mg, *TestDiet, USA*) that were placed in the magazine. The volume of water consumed by each rat over this 60-min session was measured in order to determine the amount of water each individual drank while eating 60 45 mg pellets.

Rats were then tested in one 60-min magazine training session during which 60 food pellets were delivered under a random time schedule (RT-60 seconds) over 60 min. This magazine training provided the baseline level of homeostatic water intake over one hour.

#### *SIP water*

The SIP procedure was based on a fixed-time 60-second schedule of food delivery, previously shown to induce adjunctive drinking behavior with robust and persistent inter-individual differences in its magnitude (20, 22, 23).

Twenty-four hours after the baseline session, rats underwent 21 of these FT-60s SIP sessions (20, 24). 300ml bottles filled with fresh water were weighed and inserted into the operant box immediately prior to the initiation of each session. House lights were switched on at the beginning and switched off at the end of each session. The total amount of water consumed (ml) was calculated daily as the difference between the weights of the bottle before and after the session. Individuals whose average water consumption over the last three days of training was in the upper and lower quartiles of the population were considered as high (HD, n=12) and low drinkers (LD, n=12), respectively (see 20, 21).

#### *SIP alcohol*

Twenty-four hours after the last SIP session for water, the water was replaced by 10% ethanol (*Sigma-Aldrich*). As predicted, some individuals only acquired SIP when the adjunctive response was directed to alcohol, and we therefore stratified rats based on their average alcohol consumption over the last three sessions of SIP with alcohol as “high drinker water-high drinker alcohol” (HDw-HDa, n=6) or “high drinker water-low drinker alcohol” (HDw-LDa, n=6), “low drinker water-high drinker alcohol (LDw-HDa, n=6) or “low drinker water-low drinker alcohol” (LDw-LDa, n=6). For clarification purposes, rats from the groups HDw-LDa and LDw-HDa were labelled Water Copers (WC) and Alcohol Copers (AC) rats respectively in the main manuscript.

#### *Resistance to adulteration*

Rats underwent a session of SIP with alcohol as described above, but the 10% ethanol solution was supplemented with 0.1g/L quinine (*Sigma-Aldrich*) (25). The resistance to adulteration was calculated as the ratio between alcohol+quinine intake over the average alcohol consumption over the last three sessions of SIP alcohol.

### Blood alcohol level quantification

Blood samples were collected immediately after the last session of SIP with alcohol. Rats were anaesthetized under isoflurane and blood (0.5-1mL) was collected from the sublingual vein in heparinized tubes. After centrifugation (3000rpm, 5min, 4ºC), the plasma was collected and the blood alcohol levels (BAL) were determined in triplicate in mg/ml using Analox AM1 (26).

Supplementary Statistics

### Statistical analyses

Data are presented as number of individuals, mean (for some radar plots) or mean +/- standard error of mean (SEM). Assumptions for parametric analyses, namely homogeneity of variance, sphericity and normality of distribution were verified prior to each analysis with Cochran, Mauchly and Shapiro-Wilk’s tests respectively. Behavioral data were subjected to repeated measure, one-, two- factorial analyses of variance (ANOVAs) or non-parametric analyses, i.e. Kruskal-Wallis or Mann-Whitney U tests when the datasets did not meet assumptions for parametric analyses. The experimental groups (LR/HR, HA/LA, GT/ST, LSP/HSP, LNP/HNP, SE/EE, number of positive addiction-like criteria…) were used as between-subject factor and, depending on the analysis, the arm (open/closed arms) and the time (number of sessions or session blocks) were used as within-subject factors. Chi-squared test (maximum likelihood) was used to test the assumption of independent normally distributed data wherever necessary. Upon confirmation of significant main effects differences among individual means were further analyzed using the Newman-Keuls post-hoc test and planned comparisons wherever appropriate. The significance level was set at p ≤ 0.05 and for all significant analyses, the effect size was reported by the partial eta-squared value (ηp^2^).

### Experiment 3

EE and SE rats displayed a similar pattern of alcohol intake over the period of exposure to the unpredictable two-bottle choice of alcohol and water [main effect of group: F_1,22_<1; session: F_19,148_=30.816, p<0.001; and group x session interaction: F_19,148_<1)].

At day 225, two groups in each environmental condition were segregated so as to not differ in their alcohol intake both over training pre-abstinence and during relapse [EE, training pre-abstinence: main effect of group: F_1,10_<1, session: F_46,460_=20.081, p<0.001; group x session interaction: F_46,460<_1, relapse: Mann-Whitney, U=18, p>0.05], [SE, training pre-abstinence: main effect of group: F_1,10_<1, session: F_46,460_=10.769, p<0.001; group x session interaction: F_46,460_<1, relapse: Mann-Whitney, U=13, p>0.05]. One group of each housing condition was subsequently exposed to alcohol adulteration as presented in the main text.

### Experiment 4

Rats from each group (HDw-HDa, HDw-LDa, LDw-HDa and LDw-LDa) displayed similar average levels of alcohol intake during the last 3 post-adulteration sessions of SIP alcohol (D60-62) as compared to the last 3 pre-adulteration sessions of SIP alcohol (D54-56). Thus, the BAL measured in post-adulteration baseline sessions was reflective of that seen in rats prior to be exposed to quinine [main effect of phenotype: F_3,20_=13.049, p<0.0001; pre/post-adulteration: F_1,20_<1 and phenotype x sessions interaction: F_3,20_=2.4203, p>0.05].

Control rats selected in the two-bottle choice procedure to match the overall alcohol intake of that displayed by those exposed to the SIP procedure, showed similar differences between subpopulations (i.e. HDw-HDa, HDw-LDa, LDw-HDa and LDw-LDa) [subpopulation x cohort interaction: F_3,38_<1].

Supplementary Figures

**Figure S1.**

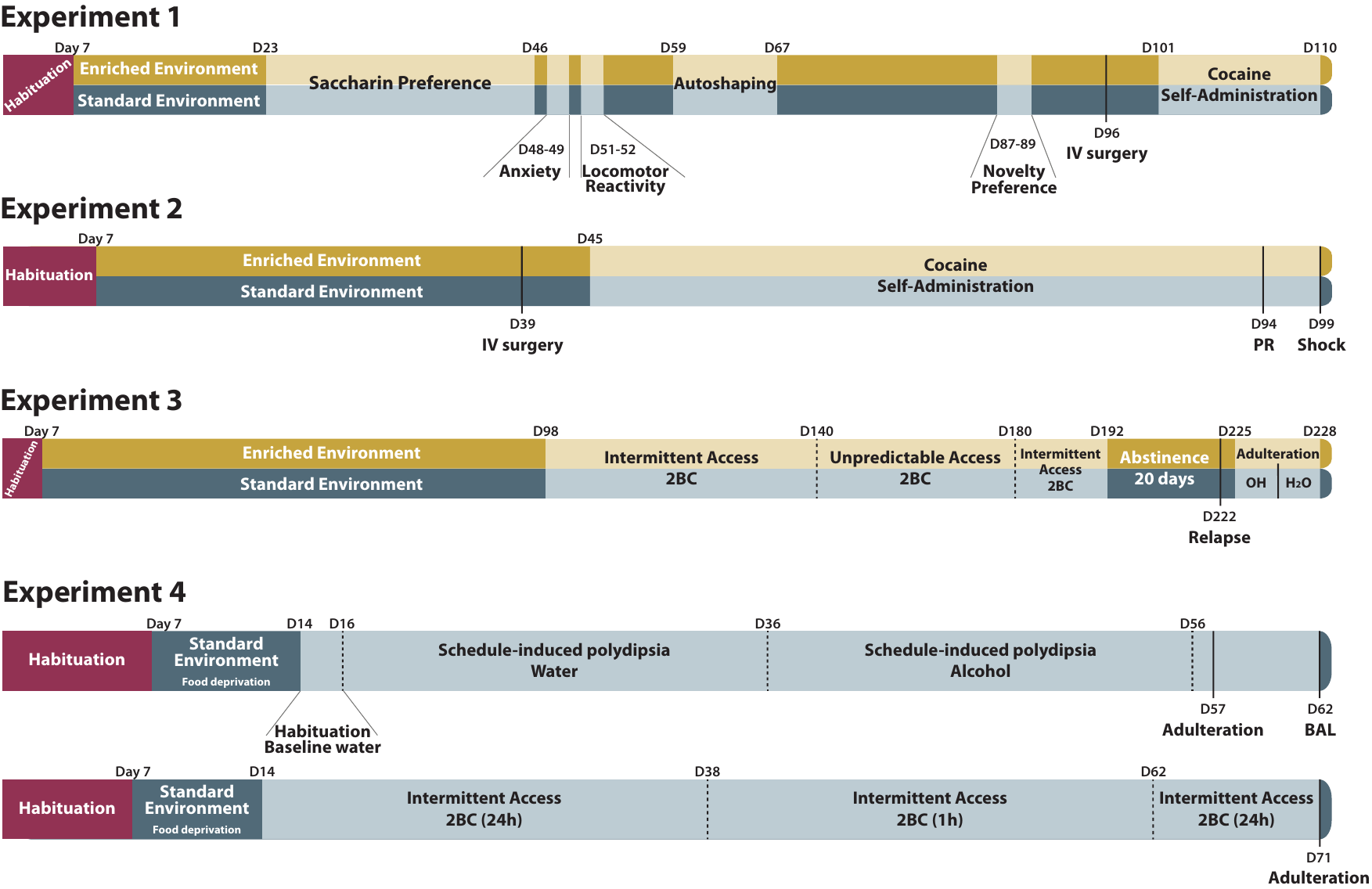

***Experimental timelines of the four experiments.***

**Experiment 1:** Following a week of habituation to the vivarium, male Sprague Dawley rats (n=48) were housed in EE (n=24) or SE (n=24) until the end of the experiment. Sensitivity to natural reward was assessed by a two-bottle choice between water and saccharine as alternative reinforcers carried out from day 23 to day 46. Rats were then screened for their anxiety-related behavior in an Elevated Plus Maze and their locomotor reactivity to novelty in an open field at days 48-49 and 51-52 respectively. Two rats from the SE group and one rat from the EE group were excluded from the results of the Elevated Plus Maze (EPM) task after falling from the maze.

The individual differences in ascribing incentive value to a CS was assessed in an autoshaping (sign-tracking) task between day 59 and day 67 and the preference for novelty over a familiar environment was tested at days 87-89 in a Novelty Place Preference (NPP) test. One rat from the EE group escaped from its NPP box and was therefore excluded from the analyses. Due to a software failure, the data of an entire NPP session (4 EE rats) could not be acquired, thus these rats were excluded from the analysis. At day 96, rats received an indwelling catheter into their right jugular vein under anaesthesia and after five days, the influence of differential housing conditions on the self-administration of cocaine (SA) was assessed. SA training started with six 3.5-hour daily sessions under a fixed ratio (FR) 1 schedule of reinforcement (FR1) followed by one under FR3 session and three under FR5.

**Experiment 2:** Following a week of habituation to the vivarium, male Sprague Dawley rats (n=48) were housed in EE (n=24) or SE (n=24) until the end of the experiment. At day 39, rats received an indwelling catheter into their right jugular vein under anaesthesia and, after five days, were trained to self-administer cocaine starting with five 3.5-hour daily sessions under FR1 followed by 2 sessions under FR3 and 40 sessions under FR5.

The individual vulnerability to develop addiction-like behavior was assessed as (i) the inability to refrain from responding despite the drug being signalled as unavailable, during so-called no-drug periods (days 92-93) ; (ii) a high motivation for the drug, assessed over a single session under a progressive-ratio schedule of reinforcement and (iii) the persistence of cocaine SA despite punishment, assessed during a single 40 min long session (day 99) that was preceded by four baseline sessions during which instrumental responses were punished response-contingently.

Twelve rats (EE group: n=5; SE group: n=7) were excluded from the final analysis due to loss of catheter patency.

**Experiment 3:** Following a week of habituation to the vivarium, male Sprague Dawley rats (n=24) were housed in EE (n=12) or SE (n=12) until the end of the experiment. On day 98, rats were exposed to 21 two-bottle choice sessions under intermittent access. Rats were then given 20 two-bottle choice sessions under unpredictable access (see methods) followed by six sessions under intermittent access. Rats were returned to their home cage for a period of 20 days abstinence, following which, on day 222 they were exposed to a four hour two-bottle choice session (relapse session) followed by a twenty-four hour two-bottle choice session. From day 225 onwards, EE and SE cohorts were both split into two subpopulations of rats displaying similar levels of drinking behavior across their alcohol exposure history (pre-abstinence training and during relapse). One subpopulation of rats was exposed to two sessions during which alcohol was adulterated with quinine (0.1 g/L) while the other subpopulation underwent the same procedure, but without quinine. Rats were then given two sessions during which they had access to a single bottle of water adulterated with quinine (0.1 g/L).

**Experiment 4:** Following a week of habituation to the vivarium, male Sprague Dawley rats (n=48) were housed in the standard environment (SE) only and were trained in schedule-induced polydipsia procedure (SIP). Following several habituation steps (see main text), rats were exposed to 20 1-hour SIP sessions with water followed by 20 1-hour SIP sessions with alcohol. The following day, the individual tendency to persist in drinking alcohol despite adulteration with quinine (0.1 g/L) was assessed after which rats underwent five additional sessions of SIP with alcohol over which they reached levels of alcohol intake that were similar to those they had shown prior to adulteration. Blood samples were collected immediately after the last session to measure their blood alcohol level (BAL). One blood sample from an individual of the Alcohol Copers group was discarded due to a technical issue.

In order to test whether persistence in drinking alcohol despite adulteration shown by rats trained under SIP procedure was dependent on the SIP context in which rats initiated their alcohol use, another cohort (n=24) was housed under similar housing conditions and were then given a two-bottle choice procedure under intermittent access for twelve 24-hour sessions, followed by twelve 1-hour sessions. Rats were then received an intermittent access two-bottle choice for eight 24-hour sessions followed by a single 1-hour session during which their individual tendency to maintain alcohol drinking despite adulteration was measured.

We then sought to verify that the persistence of drinking alcohol despite adulteration shown by rats trained under a SIP procedure was attributable to their acquisition of alcohol drinking in the self-medication context of SIP. Another cohort of twenty-four male SD rats was housed in a similar SE under food deprivation and were exposed to a two-bottle choice procedure under intermittent access for twelve 24-hour sessions, followed by twelve 1-hour sessions. Rats then received an intermittent access to two-bottle choice for eight 24-hour sessions followed by a single 1-hour session during which their individual tendency to maintain alcohol drinking despite adulteration was measured.

**Figure S2.**

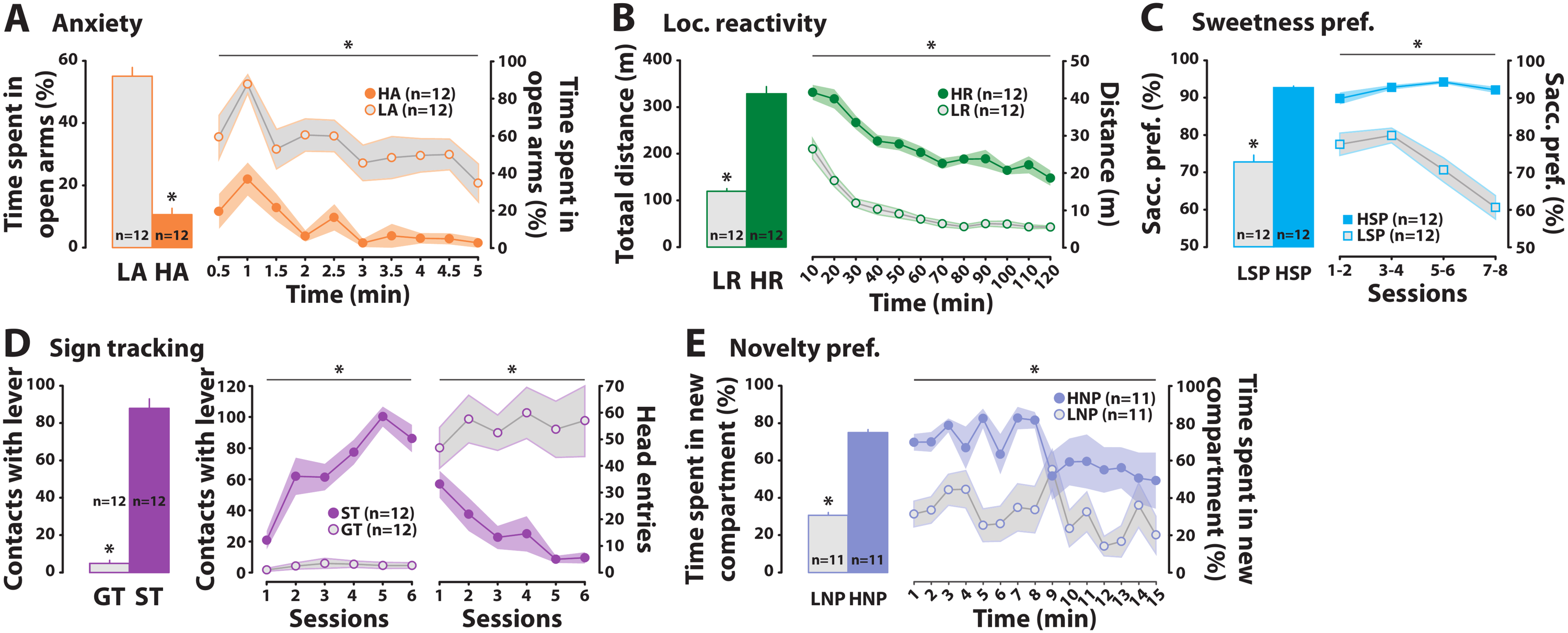

***Multidimensional behavioral profile of heterogenous rat populations: inter-individual differences in behavioral features of resilience or vulnerability to addiction are observed irrespective of differential housing conditions.***

**A)** Low anxious rats (LA) displayed a higher percentage of time spent in the open arms of the elevated plus maze than high anxious (HA) rats [main effect of group: F_1,22_=163.63, p<0.001, ηp^2^=0.88] and this difference was stable across the 5 min session. **B)** High responder (HR) rats displayed a higher reactivity to a novel environment as compared to low responder (LR) rats [main effect of group: F_1,22_=167.68, p<0.001, ηp^2^=0.88, time: F_11,242_=53.605, p<0.001, ηp^2^=0.70 and group x time interaction: F_11,242_=2.0068, p<0.05, ηp^2^=0.08]. **C)** High saccharin preferring (HSP) rats consistently drank much more saccharin than water as compared to low saccharin preferring (LSP) rats over test sessions [main effect of group: F_1,22_=117.36, p<0.001, ηp^2^=0.84] and this effect was stable across sessions. **D)** Sign tracker (ST) rats displayed a higher level of lever press on the cue-associated lever as compared to goal tracker (GT) rats over the last three training sessions [main effect of group: F_1,22_=244.05, p<0.001, ηp^2^=0.91] the average of which is presented on the left, and across sessions [main effect of group: F_1,22_=166.42, p<0.001, ηp^2^=0.88; session: F_5,110_=12.906, p<0.001, ηp^2^=0.37 and group x session interaction: F_5,110_=11.190, p<0.001, ηp^2^=0.34]. In marked contrast, GT rats displayed a higher level of head entries in the food magazine as compared to ST rats across sessions [main effect of group: F_1,22_=21.965, p<0.001, ηp^2^=0.50; session: F_5,110_=1.4035, p>0.05 and group x session interaction: F_5,110_=3.3251, p<0.01, ηp^2^=0.13]. **E)** High novelty preferring (HNP) rats displayed a higher percentage of time spent in the novel compartment when given the opportunity freely to explore both this new and the known compartment, as compared to low novelty preferring (LNP) rats [main effect of group: F_1,20_=427.51, p<0.001, ηp^2^=0.95] and this effect was stable over the 15 min. *p≤0.05.

**Figure S3.**

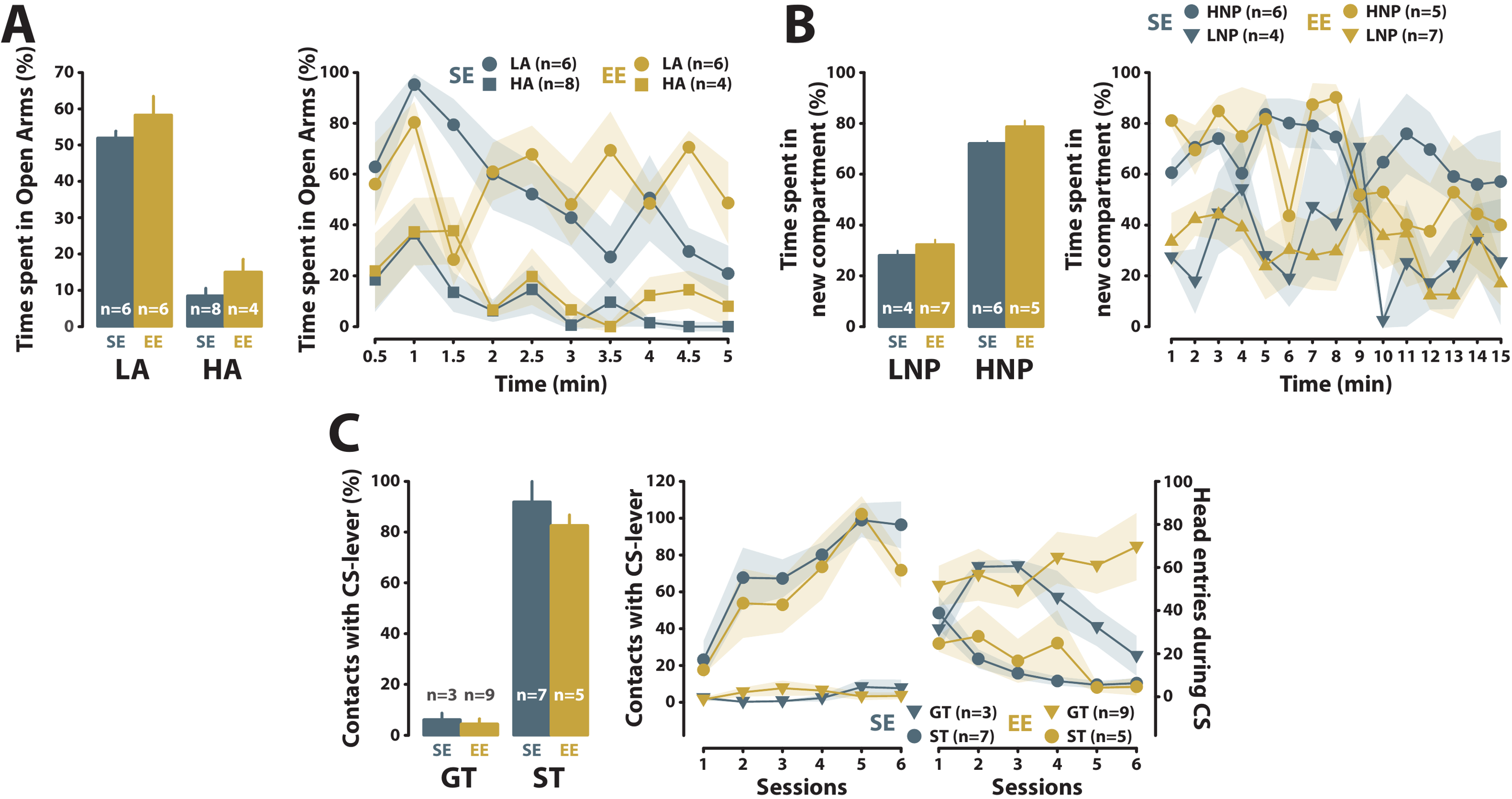

***Housing conditions have little influence on the magnitude of the behavioral manifestation of the underlying traits of vulnerability to addiction.***

**A)** High and low anxious rats from the EE did not differ from those from the SE in the percentage of time spent in the open arms of the elevated plus [environment x phenotype interaction: F_1,20_<1, and time x environment x phenotype interaction: F_9,180_=2.3923, p<0.05, ηp^2^=0.11]. **B)** Similarly, high and low novelty preference rats from SE did not differ respectively from those from an EE in the percentage of time spent in the novel compartment in a NPP task [phenotype x environment interaction: F_1,18_<1 and time x environment x phenotype interaction: F_14,252_<1]. **C)** Sign and goal tracker rats from the EE did not differ over the three last sessions from those form the SE respectively in the levels of lever press on the cue-associated lever [environment x phenotype interaction: F(1,20)<1] and across the 6 sessions [environment x phenotype interaction: F(1,20)=1.1321, p>0.05 and session x environment x sign tracking interaction: F_5,100_<1] or head entries in the food magazine [environment x phenotype interaction: F_1,20_<1 and session x environment x phenotype interaction: F_5,100_=3.4041, p<0.01, ηp^2^=0.15]. **D)** Low saccharin preferring and high saccharin preferring rats from the EE did not differ respectively from those form the SE in their preference for saccharin over water [environment x phenotype interaction: F(1,20)=1.6551, p>0.05] and was consistent over time within each session.

**Figure S4.**

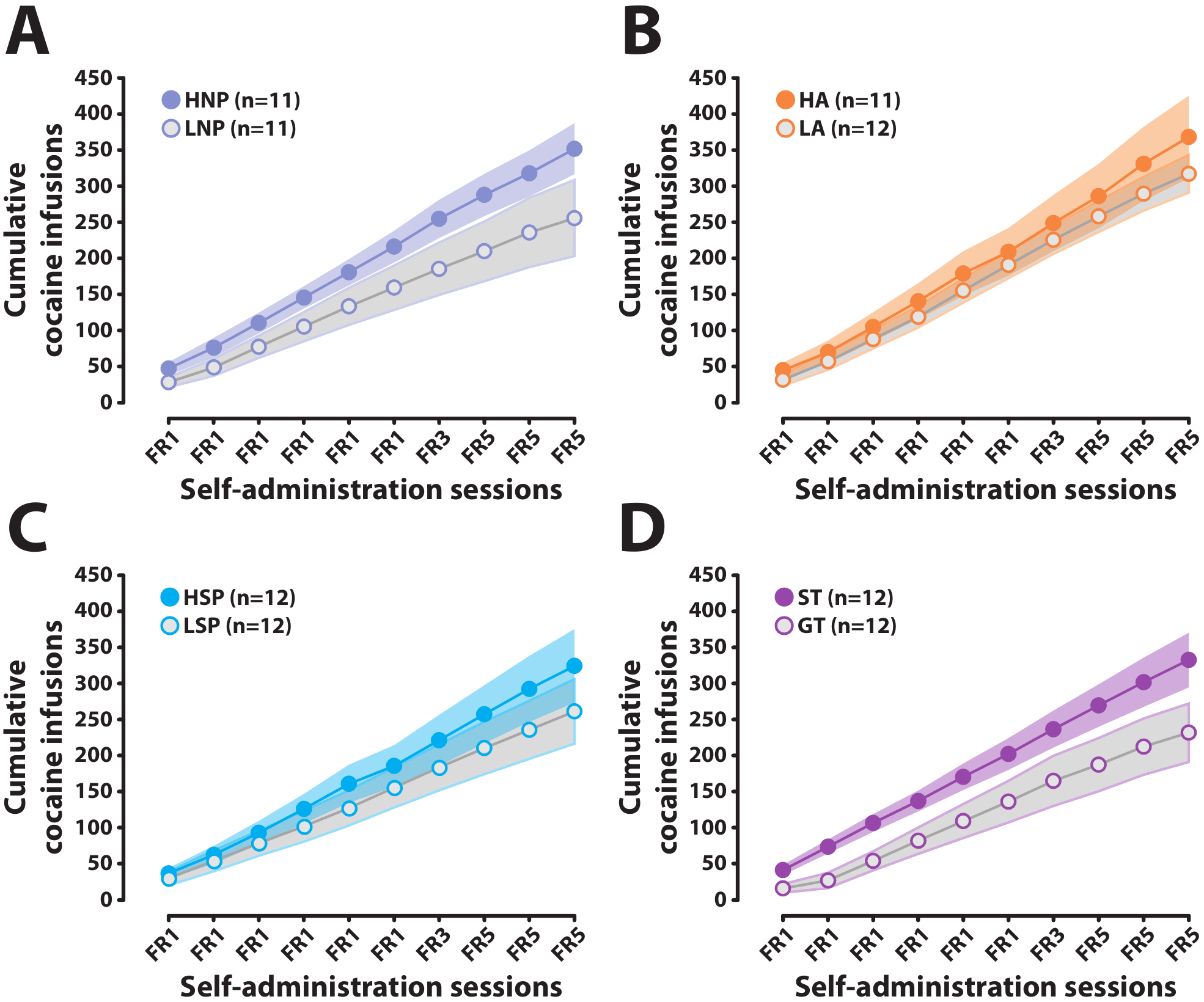

***The individual propensity to acquire cocaine SA is not predicted by novelty preference, anxiety, saccharin preference or sign tracking.***

No differences were observed over the 10 daily sessions of cocaine self-administration under increasing fixed ratio schedules (FR) of reinforcement between rats characterised as **(A)** high (HNP) or low novelty preferring (LNP) [main effect of group: F_1,20_=2.5098, p>0.05; session: F_9,180_=78.603, p<0.001, ηp^2^=0.80 and group x session interaction: F_9,180_=1.5574, p>0.05], **(B)** high anxiety (HA) or low anxiety (LA) [main effect of group: F_1,21_<1; session: F_9,189_=133.25, p<0.001, ηp^2^=0.86 and group x session interaction: F_9,189_<1], **(C)** high (HSP) or low saccharine preference (LSP) [main effect of group: F_1,22_<1; session: F_9,198_=70.076, p<0.001, ηp^2^=0.76 and group x session interaction: F_9,180_<1]. (**D)** Sign trackers (ST) or goal trackers (GT), although a trend was observed [main effect of group: F_1,22_=4.2827, p=0.051; session: F_9,198_=83.348, p<0.001, ηp^2^=0.79 and group x session interaction: F_9,180_=1.3188, p>0.05].

**Figure S5.**

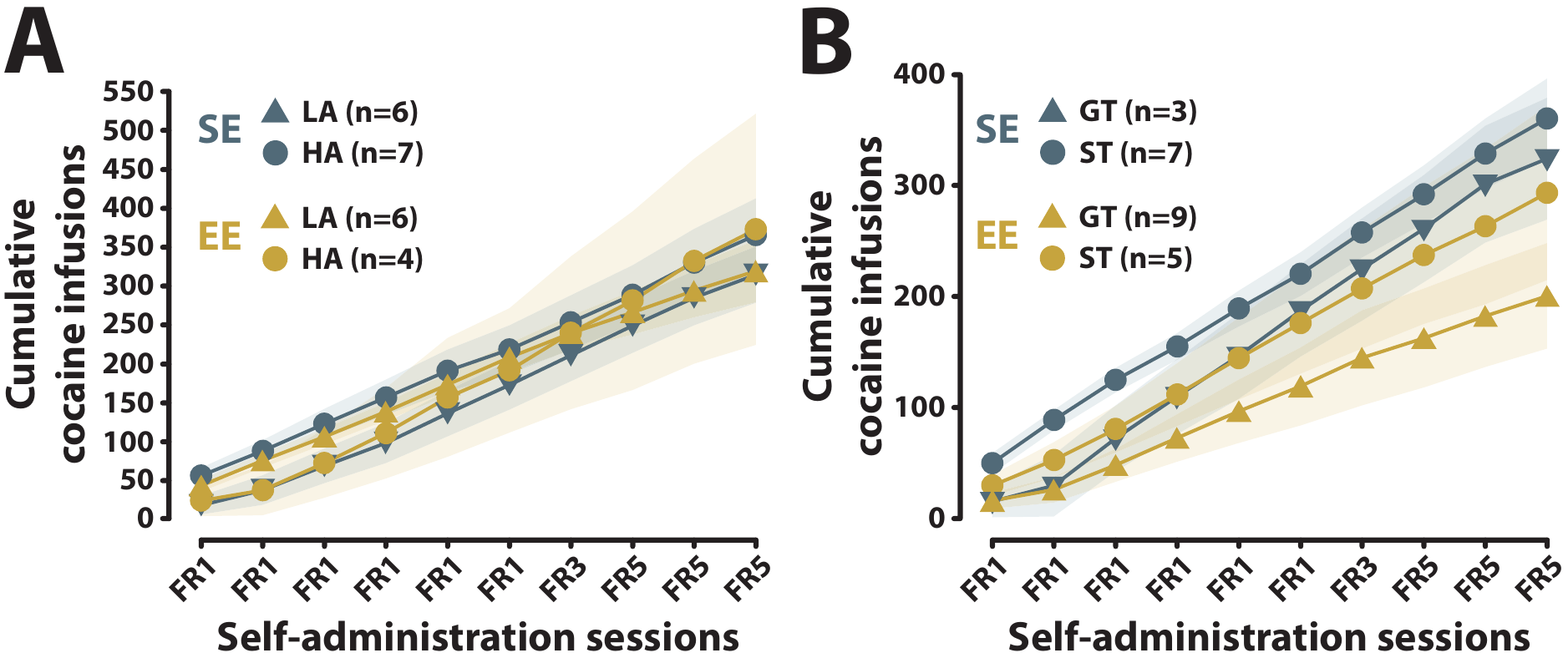

***Housing conditions had no influence on the role of anxiety and sign/goal tracking in the propensity to acquire cocaine SA.***

High anxiety (HA) and Low anxiety (LA) rats **(A)** as well Sign and Goal trackers **(B)** displayed the same propensity to acquire cocaine SA irrespective of their housing conditions [HA/LA: environment x phenotype interaction: F_1,19_<1 and session x environment x phenotype interaction: F_9,171_<1, ST/GT: environment x phenotype interaction: F_1,20_<1; session x environment x phenotype interaction: F_9,180_<1].

**Figure S6.**

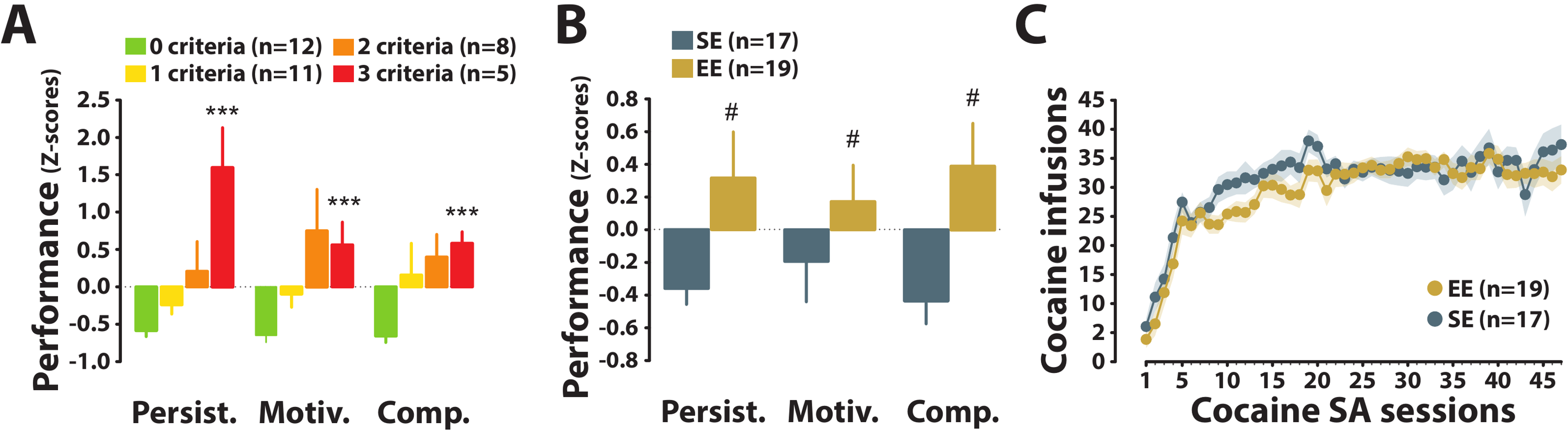

***The increased propensity to develop addiction-like behaviour associated with environmental enrichment was not due to a differential history of cocaine self-administration.***

**A)** The criterion-dependent addiction severity shown by 0, 1, 2 and 3-crit rats, regardless of their housing conditions (figure 3A), reflected criterion-dependent differences in each of the three addiction-like behaviours, namely the inability to refrain from drug seeking when the drug is signalled as not available (persistence -Persist.-), their higher motivation for the drug (Motiv.) measured under a task-specific progressive ratio schedule of reinforcement and their persistence in self-administering cocaine despite punishment (e.g., compulsive drug SA -Comp.-) [main effect of group: F_3,32_=17.97, p<0.001, ηp^2^=0.63, test: F_2,6_<1 and group x test interaction: F_6,64_=1.23, p=0.3]. Consequently, 3crit rats displayed significantly higher scores than 0crit rats for each of the behaviours (***: p<0.001). **B)** Similarly, the influence of the different housing conditions on addiction severity (Figure 3B) is reflected in the higher scores of EE as compared to SE rats for each of the addiction-like behaviours [main effect of environment: F_1,34_=8.82, p<0.01, ηp^2^=0.21, behaviour: F_1,2_<1 and environment x test interaction: F_1,68_<1] (#: p<0.01). **C)** These differences between housing conditions in the propensity to develop addiction-like behaviour were not attributable to a difference in cocaine intake over their drug taking history. Even though EE rats were initially slower than SE rats to acquire cocaine SA [main effect of environment x session interaction: F_1,46_=1.44, p=0.03], the two groups did not differ overall in their daily cocaine intake over the course of the 47 daily sessions that preceded the assessment of addiction-like behaviour [main effect of environment: F_1,34_<1].

**Figure S7.**

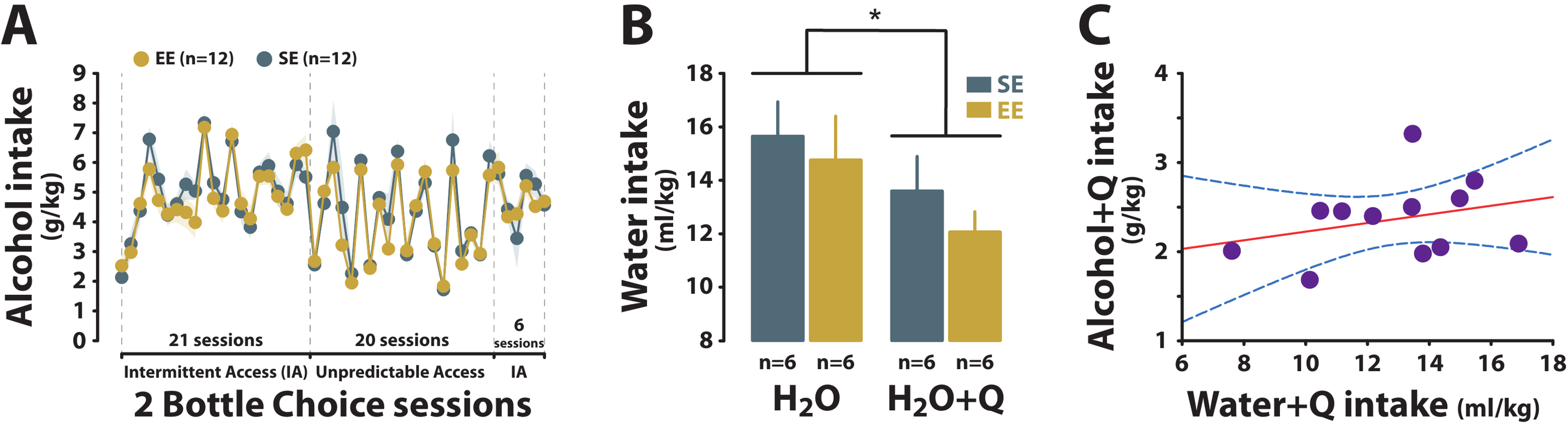

***Figure S7: Housing conditions had no differential influence on alcohol intake in a two bottle-choice procedure or on the aversiveness of quinine.***

A) EE and SE rats did not differ in their daily intake of alcohol over the 47 daily sessions of intermittent or unpredictable access to alcohol in a two bottle choice procedure that preceded the assessment of relapse and the persistence in drinking alcohol that had been adulterated with quinine [main effect of environment: F_1,22_<1; session: F_46,1012_=27.27, p<0.01, ηp^2^=0.55 and environment x session interaction: F_46,1012_<1]. **B)** EE and SE did not differ in the degree of decrease in water intake when it was supplemented with 0.1g/L quinine [main effect of environment: F_1,10_<1; treatment: F_1,10_=12.01, p<0.01, ηp^2^=0.55 and environment x treatment interaction: F_1,10_<1]. **C)** The intake of adulterated alcohol did not correlate with that of adulterated water irrespective of the housing conditions [R=0.29, p>0.05]. H_2_0: water, H_2_O+Q: water + quinine.

**Figure S8.**

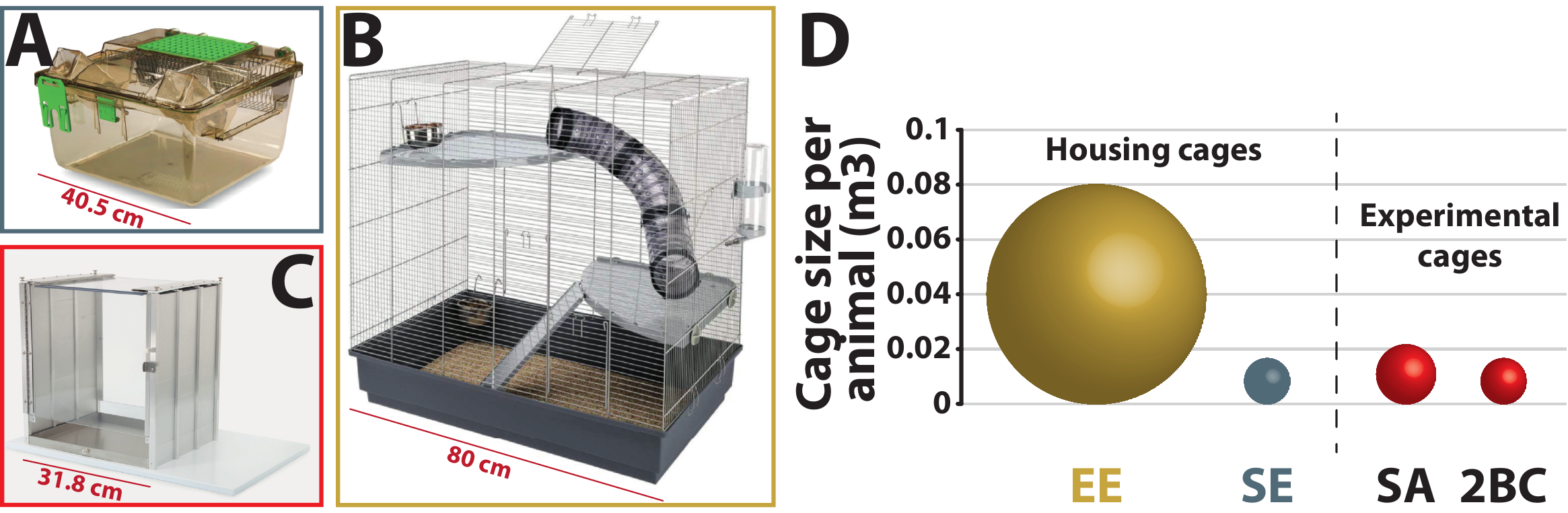

***The drug self-administration setting is impoverished as compared to the enriched housing environment.***

Rats housed in SE cages (40.5x35.5x23cm) **(A)** or EE cages (80x50x80cm) **(B)** acquired cocaine self-administration (SA) in operant chambers (31.8 cm x 25.4 cm x 26.7 cm) **(C)** or alcohol SA in the SE cages. **(D)** In addition to being housed with 3 other rats and having access to toys, wheel, platforms and hammocks (as previously described), EE rats benefit in their housing environment from an individual space almost 4 times greater than that experienced in the drug SA settings. This dramatic negative contrast between housing and testing conditions experienced by EE rats is different to that experienced by SE rats for whom the housing conditions provide a rather individual space similar to, or smaller than, that of the drug SA settings.

**Figure S9.**

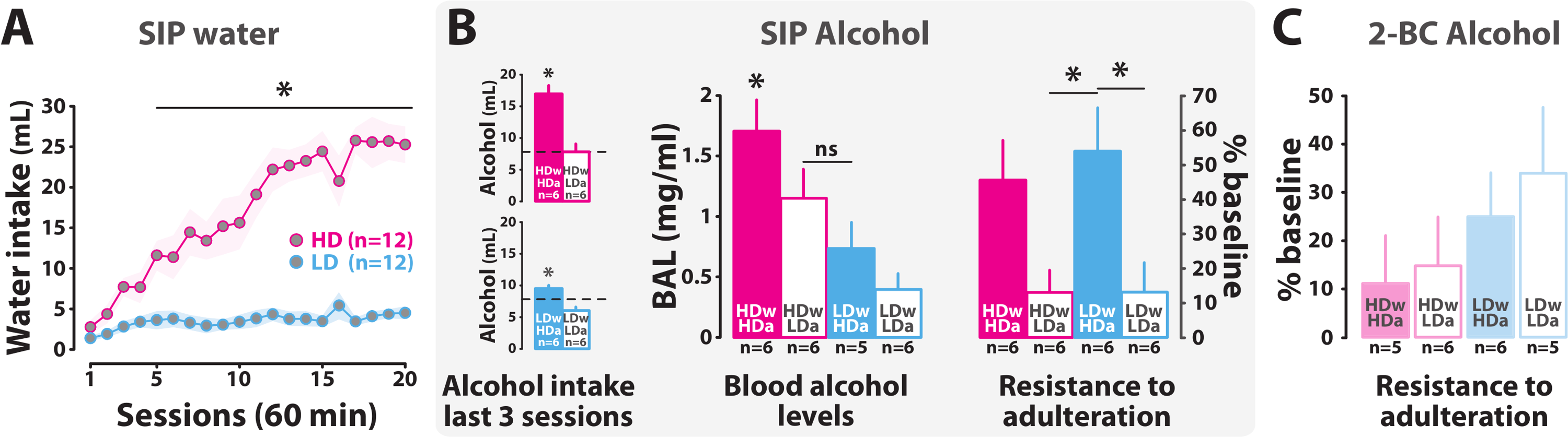

***Figure S9: Compulsive alcohol drinking is specific to the self-medication context.***

**A)** the upper and lower quartiles (n=12 each) of the SE population exposed to the SIP procedure were characterised as high drinker (HD) and low drinker rats (LD) respectively, based on the average water consumption over the last 3 sessions of SIP with water (20, 21) so that HD rats developed SIP-associated increases in water intake from day 5 onwards whereas LD rats never increased their water intake [main effect of phenotype: F_1,22_=53.975, p<0.001, ηp^2^=0.71; session: F_19,418_=20.133, p<0.001, ηp^2^=0.48 and phenotype x session interaction: F_19,418_=14.236, p<0.001, ηp^2^=0.39]. **B)** When alcohol was introduced in place of water in the SIP procedure, rats belonging to the HD and LD groups differed in their drinking alcohol levels. Thus, half of the HD rats persisted in drinking high amounts of alcohol (HDwHDa, n=6) while others drank substantially less alcohol than water (HDwLDa, n=6) so that HDwHDa differed from HDwLDa [main effect of phenotype: F_1,10_=24.474, p<0.001, ηp^2^=0.71]. Similarly, some LD rats eventually developed high levels of alcohol intake, thereby showing the ability to acquire a coping strategy with alcohol only (LDwHDa, n=6). The other LD rats did not show much more interest in alcohol than they did in water under SIP (LDwLDa, n=6). Consequently, LDwHDa rats drank significantly more alcohol during the last 3 sessions of SIP with alcohol than LDwLDa rats [main effect of phenotype: F_1,10_=21.206, p<0.001, ηp^2^=0.68].

Whereas HDwLDa and LDwHDa rats did not differ in their overall alcohol intake [main effect of phenotype: F_1,10_=1.2771, p>0.05, data not shown], or in their acute blood alcohol [main effect of phenotype: F_3,19_=6.8625, p<0.01, ηp^2^=0.52], they were significantly different with regards to compulsivity, in that only the latter persisted in drinking alcohol despite adulteration [main effect of phenotype: F_3,20_=4.5419, p<0.05, ηp^2^=0.41]. **C)** Rats that acquired alcohol in a standard 2 bottle choice procedure in their home cage and matched with regards to their alcohol intake with the various SIP-alcohol related groups (i.e. HDwHDa, HDwLDa, LDwHDa and LDwLDa), did not differ in their sensitivity to alcohol adulteration [Kruskal-Wallis: H=3.8347, p>0.05].
